# Targeted molecular profiling of rare cell populations identifies olfactory sensory neuron fate and wiring determinants

**DOI:** 10.1101/2020.07.08.191528

**Authors:** J. Roman Arguello, Liliane Abuin, Jan Armida, Kaan Mika, Phing Chian Chai, Richard Benton

## Abstract

Determining the molecular properties of neurons is essential to understand their development, function and evolution. We used Targeted DamID (TaDa) to characterize RNA polymerase II occupancy and chromatin accessibility in selected Ionotropic Receptor (IR)-expressing sensory neurons in the *Drosophila* antenna. Although individual populations represent a minute fraction of cells, TaDa is sufficiently sensitive and specific to identify the expected receptor genes. Unique *Ir* expression is not linked to substantial differences in chromatin accessibility, but rather to distinct transcription factor profiles. Heterogeneously-expressed genes across populations are enriched for neurodevelopmental factors, and we identify functions for the POU-domain protein Pdm3 as a genetic switch of Ir neuron fate, and the atypical cadherin Flamingo in segregation of neurons into discrete glomeruli. Together this study reveals the effectiveness of TaDa in profiling rare neural populations, identifies new roles for a transcription factor and a neuronal guidance molecule, and provides valuable datasets for future exploration.

## Introduction

Nervous systems are composed of vast numbers of neuron classes with specific structural and functional characteristics. Determining the molecular basis of these properties is essential to understand their emergence during development and contribution to neural circuit activity, as well as appreciating how neurons exhibit plasticity over both short and evolutionary timescales.

The olfactory system of adult *Drosophila melanogaster* is a powerful model to identify and link molecular features of neurons to their anatomy and physiology. The *Drosophila* nose (antenna) contains ∼50 classes of olfactory sensory neurons (OSNs), which develop from sensory organ precursors cells specified in the larval antennal imaginal disc^1-3^. Each OSN is characterized by the expression of a unique olfactory receptor (or, occasionally, receptors), of the Odorant Receptor (OR) or Ionotropic Receptor (IR) families. These proteins function – together with broadly-expressed, family-specific co-receptors – to define neuronal odor-response properties^4-7^. Moreover, the axons of all neurons expressing the same receptor(s) converge to a discrete and stereotyped glomerulus within the primary olfactory center (antennal lobe) in the brain, where they synapse with both local interneurons (LNs) and projection neurons (PNs)^3,8,9^. While the unique properties of individual OSN classes are assumed to reflect distinct patterns of gene expression, it is unclear how many molecular differences exist between sensory neuron populations, which types of genes (beyond receptors) distinguish OSN classes, and whether gene expression differences reflect population-specific chromatin architecture.

Exploration of the molecular properties of specific neuron types has been revolutionized by single-cell RNA sequencing (scRNA-seq) technologies^10^. However, the peripheral olfactory system of *Drosophila* poses several challenges to single-cell isolation: OSNs are very small (∼2-3 μm soma diameter) and up to four neurons are tightly embedded under sensory sensilla (a porous cuticular hair that houses the neurons’ dendrites) along with several support cells that seal each neuronal cluster from neighboring sensilla^11,12^. Furthermore, each OSN class represents only a very small fraction of cells of the whole antenna, as few as 5-10 neurons of a total of ∼3000 neurons and non-neuronal cells^11^. Recent work^13^ – discussed below – successfully profiled RNA levels in OSNs at a mid-developmental time-point (42-48 h after puparium formation (APF)), before these cells become encased in the mature cuticle.

To characterize the molecular profile of specific populations of OSNs we sought to use the Targeted-DNA adenine methyltransferase identification (TaDa) method^14^. TaDa entails expression of an RNA polymerase II subunit (Rp215, hereafter PolII) fused to the *E. coli* DNA adenine methyltransferase (Dam). This complex (or, as control, Dam alone) is expressed within a specific cell population (“targeted”) under the regulation of a cell type-specific promoter using the Gal4/UAS system. When bound to DNA, Dam:PolII methylates adenine within GATC motifs; enrichment of methylation at genes compared to free Dam control samples provides a measure of RNA PolII occupancy, which is correlated with genes’ transcriptional activity^14^ (Figure 1A). In addition to indicating RNA PolII occupancy within a given cell type, TaDa datasets can also provide information on chromatin accessibility (CATaDa)^15^ (Figure 1A). Analysis of the methylation patterns across the genome generated by untethered Dam is assumed to reflect the openness of chromatin, analogous to – and in good agreement with – ATAC-seq datasets^15,16^.

**Figure 1.**
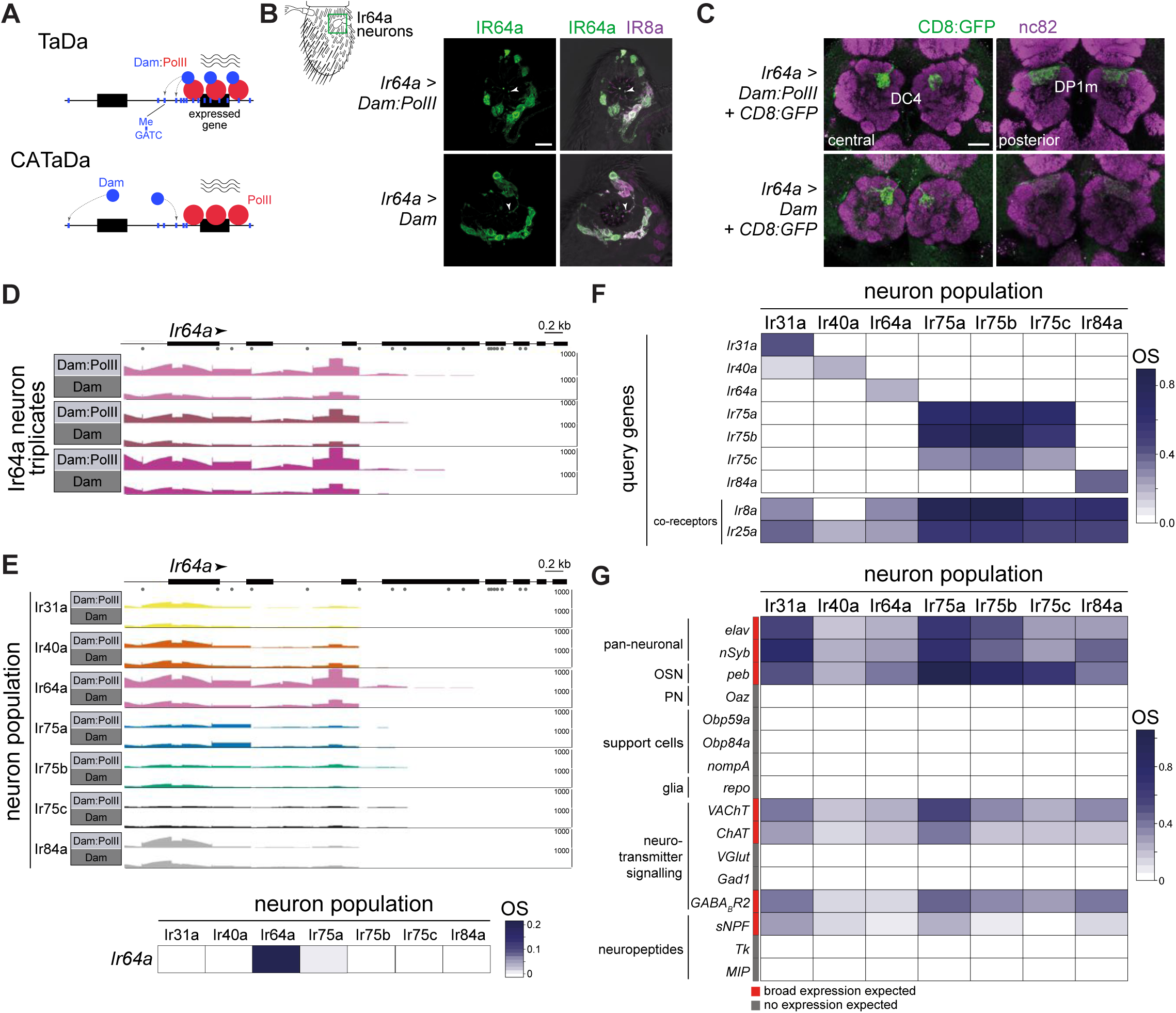
Targeted DamID of OSN populations. (A) Schematic of the principles of Targeted DamID (TaDa) and Chromatin accessibility TaDa (CATaDa). TaDa reports on the enrichment of GATC methylation (Me; blue lines) by an RNA PolII:Dam fusion (top) relative to Dam alone (bottom), thereby identifying PolII-transcribed genes. In CATaDa, analysis of methylation patterns of free Dam provides information on chromatin accessibility. (B) Immunofluorescence for IR64a and IR8a on antennal sections of animals expressing in Ir64a neurons either Dam:PolII (*Ir64a-Gal4/+*;*UAS-LT3-Dam:Rp215/+*) or Dam alone (*Ir64a-Gal4/+;UAS-LT3-Dam/+*). The approximate field of view shown is indicated by the green box on the antennal schematic on the left. The merged fluorescent channels are overlaid on a brightfield background to reveal anatomical landmarks. The arrowheads point to IR64a that is localized in the neuron sensory endings with the sensillar hairs. Scale bar = 10 µm. (C) Immunofluorescence for GFP and the neuropil marker nc82 on whole mount brains of animals expressing in Ir64a neurons a CD8:GFP reporter together with Dam:PolII (*Ir64a-Gal4/+;UAS-LT3-Dam:Rp215/UAS-mCD8:GFP*) or free Dam (*Ir64a-Gal4/+;UAS-LT3-Dam/UAS-mCD8:GFP*). Two focal planes are shown to reveal the two glomeruli (DC4 and DP1m) innervated by Ir64a neurons. Background fluorescence in the GFP channel was slightly higher across the brain in the Dam-alone samples. Scale bar = 20 µm. (D) Quantification of sequence reads of methylated GATC fragments mapping to the *Ir64a* gene (transcribed left-to-right, as indicated by the arrowhead; gray dots under the gene model indicate the approximate position of GATC motifs). The transcription start site of *Ir64a* is unknown. For each of three replicate experiments, the top row is the Dam:PolII sample, which has higher read counts compared to the Dam-alone sample below. Numbers on the right of each row indicate the read depth. The OS is based on data across the full gene body, and the FDR takes into consideration the number of GATC fragments. The range of the y-axis is 0-1000 for all rows. (E) Top: comparisons among the seven Ir neuron datasets of the sequence reads mapping to the *Ir64a* gene for the Dam:PolII and Dam samples (a single representative replicate is shown for each). Genotypes are of the form shown in (B), except for Ir75a and Ir75c neuron populations where third chromosome *Ir-Gal4* driver transgenes were used. Only the Ir64a population displays a higher read count for Dam:PolII samples compared to Dam-alone samples. The range of the y-axis is 0-1000 for all rows. Bottom: heatmap representation of the occupancy scores (OSs; color scale on the right) for *Ir64a* calculated across triplicates for each of the seven neuron populations (see Methods). (F) OS heatmap for the seven target *Ir* genes, as well as the main *Ir* co-receptor genes, in the seven Ir neuron populations. OS and FDR values associated with each *Ir* gene within the corresponding neuron population are shown in Table S1. (G) OS heatmap of diverse positive- and negative-control genes in the seven Ir neuron populations (i.e., with known or expected expression/lack of expression). See text for definitions of abbreviations.

The selectivity of expression of Dam:PolII or Dam avoids the requirement for cell isolation prior to genomic DNA extraction. Previous applications of TaDa in *Drosophila* have profiled relatively abundant cell populations in central brain tissue, the intestine or the male germline^14,17-19^. Although OSN classes can be effectively labeled using transgenes containing promoters for the corresponding receptor genes, it was unclear whether TaDa would be sufficiently sensitive to measure transcriptional activity from an individual population of sensory neurons within the antenna. We therefore set out to profile several classes of IR-expressing OSNs to test the approach and identify novel factors that distinguish the properties of these evolutionarily closely-related neuronal subtypes.

## Results

### Targeted DamID of OSN populations

To test the feasibility of TaDa for profiling *Drosophila* OSNs, we first focused on the Ir64a population, which comprises ∼16 neurons that are housed in sensilla in the sacculus (a chamber within the antenna), where they detect acidic odors^20^. We first verified that *Ir64a promoter-Gal4* (hereafter, *Ir64a-Gal4*) induction of Dam or Dam:PolII did not impact the expression or localization of the IR64a receptor (Figure 1B) or the projection of these neurons to the antennal lobe (Figure 1C). We proceeded to collect triplicate samples of ∼2000 adult antennae and processed these for TaDa (see Methods). This experimental design is expected to integrate PolII occupancy patterns from mid-pupae (when *Ir*-Gal4 expression initiates) to mature OSNs. As a positive control, we first examined the mapping of reads of methylated DNA fragments to the *Ir64a* gene. Dam:PolII consistently accessed this gene region at higher levels than Dam alone, particularly near the 5’ end of the locus (Figure 1D), consistent with previously-described genome-wide PolII occupancy patterns^14^.

We extended the TaDa approach using drivers to target six additional individual OSN classes, including all acid-sensing Ir populations (Ir31a, Ir75a, Ir75b, Ir75c, Ir84a)^21,22^, as well as the hygrosensory Ir40a neurons^23,24^; these populations range from ∼8 to ∼25 cells per antenna^25^. As expected, the enrichment of Dam:PolII access to the *Ir64a* gene was specific to the Ir64a neurons (Figure 1E). We quantified “PollI occupancy scores” (OSs) by calculating the log_2_ ratio of the (normalized) number of reads from the Dam:PolII samples compared to the (normalized) number of reads in control Dam samples across the entire gene body. In Ir64a neurons, the *Ir64a* gene OS was significantly positive (OS = 0.20; FDR = 4.26e-10), while it was not significantly different from zero in the other six neuron populations (Figure 1E, bottom).

Similarly, OSs for other *Ir* genes were significantly positive in the corresponding neuronal population, but not in those in which the receptors are not expressed (Figure 1F). We noted two exceptions: first, in Ir31a neurons, *Ir40a* had a positive OS (0.12; FDR = 1.93e-13), although it was much lower than that of *Ir31a* (OS = 0.41; FDR = 2.67e-6) (Figure 1F). Second, the *Ir75a, Ir75b* and *Ir75c* genes had positive OSs in all three of the corresponding neuron populations (Figure 1F). This latter result is most likely due to these tandemly-clustered genes having overlapping transcription units^25^.

To further examine the specificity of TaDa in reporting on neuron population-specific gene expression, we plotted OSs for all members of the *Ir* repertoire, as well as *Or* and *Gustatory receptor* (*Gr*) families. The vast majority of these genes are expressed only in other antennal sensory neuron populations, distinct chemosensory organs and/or life stages^26-28^. Concordantly, only very few genes show significant OSs in any of the seven populations of Ir neurons analyzed by TaDa (Figure 1F and Figure S1). Two prominent examples are *Ir8a* and *Ir25a*, which encode co-receptors for subsets of tuning IRs^29^. *Ir8a* has a significant OS in all Ir populations except for Ir40a neurons, which is consistent with the selective expression and function of IR8a in acid-sensing OSNs but not hygrosensory neurons^29^ (Figure 1F). IR25a is broadly-expressed in most antennal neurons^29,30^ – although it may have a role in only a subset of these – which is reflected in a significant OS for *Ir25a* in all seven populations (Figure 1F). IR40a is co-expressed with another receptor, IR93a^23,24^, but we did not detect a significant OS for *Ir93a* in these neurons (Figure S1 and Table S1). This negative result may reflect the observation that while IR93a protein is readily detected^23^, *Ir93a* transcripts levels (and so potentially PolII occupancy) are extremely low^30,31^.

Of the other chemosensory receptors displaying unexpected significant OSs in these Ir neurons, many of these genes are located in the introns of other genes (e.g., *Ir47a*, within the *slowpoke 2* locus, which encodes a nervous-system expressed potassium channel^32^) or have overlapping transcripts with other genes (e.g., *Gr2a*, which overlaps with *futsch*, a gene encoding a neuronal microtubule-binding protein^33^) (Figure S1). In these cases, calculation of a specific OS for the chemosensory genes is impossible, and we suspect that in many cases the OS reflects transcription of the non-receptor gene.

Olfactory receptors have lower transcript levels in the antenna compared to other classes of genes with broader neuronal or non-neuronal expression patterns^31,34^. We extended the examination of the TaDa datasets to other genes that are known to be expressed (or not expressed) in the analyzed Ir neuron populations (Figure 1G). Two broadly-expressed neuronal genes, *elav* and *nSyb*, display significant OSs across populations, as does *pebbled* (*peb*), a molecular marker for OSNs^35^. By contrast, *oaz*, a broad marker of PNs^36^, is not significantly occupied. Many of the most highly expressed genes in the antenna are those encoding Odorant Binding Proteins (OBPs), which are transcribed in non-neuronal cells, often at >10-fold higher levels than olfactory receptor genes, as determined by bulk RNA-seq^31,34^. Two of these, *Obp59a* and *Obp84a* are present in sensilla housing Ir neurons^37,38^, but neither of these genes display significant OSs in any neuron population (Figure 1G). Similarly, markers for other support cell types, such as *nompA* (thecogen cell)^37,39^, and the glial marker *repo* are not significantly occupied (Figure 1G).

Electrophysiological experiments indicate that antennal OSNs are cholinergic^40^. In line with this functional property, genes encoding the vesicular acetylcholine transporter (*VAChT*) and choline acetyltransferase (*ChAT*) have significant OSs in all populations, while glutamatergic and GABAergic neuron markers (vesicular glutamate transporter (*VGlut*) and glutamic acid decarboxylase 1 (*Gad1*)) do not (Figure 1G). We note, however, that a GABA receptor subunit (encoded by *GABA-B-R2*) – which is heterogeneously-expressed and mediates population-specific presynaptic gain control in different Or OSNs^41^ – displays significant, but variable, OSs in different Ir populations (Figure 1G). Finally, of the multiple neuropeptides detected in the antennal lobe, only one, short neuropeptide F (sNPF), originates from OSNs^42^, where it has a role in glomerular-specific presynaptic facilitation^43^. Consistently, we observe significant and heterogeneous OSs for *sNPF*, but not other neuropeptide precursor genes that are expressed in central neuron populations (e.g., Tachykinin (*Tk*), Myoinhibiting peptide (*Mip*))^42^.

Together, these results indicate that TaDa is sufficiently sensitive and specific to detect RNA PolII occupancy at transcriptionally-active genes in very small populations of neurons embedded within the cellularly-complex antennal tissue.

### Global analysis of TaDa datasets

We next surveyed OSs at a genome-wide level, to gain insights into patterns of transcriptional activity in different Ir neuron populations. The number of genes significantly occupied by PolII ranged from 3603 to 4775 across the seven datasets, representing 20-27% of all genes (Figure 2A; Dataset S1). To investigate patterns of similarity in global OSs among these populations, we carried out hierarchical clustering of the mean OSs for all genes in the genome (Figure 2B). This analysis revealed an interesting correspondence in the clustering of OSs with the phylogeny of the corresponding receptor genes^25^: neurons expressing the most recently-duplicated receptor genes, Ir75b and Ir75c, were clustered together, with the next most similar neuron population, Ir75a, expressing the likely ancestral member of this tandem gene array^25^. Ir40a and Ir64a neurons clustered together away from the other acid-sensing populations, possibly reflecting the development of these neurons within the sacculus, rather than in sensilla located on the antennal surface.

**Figure 2.**
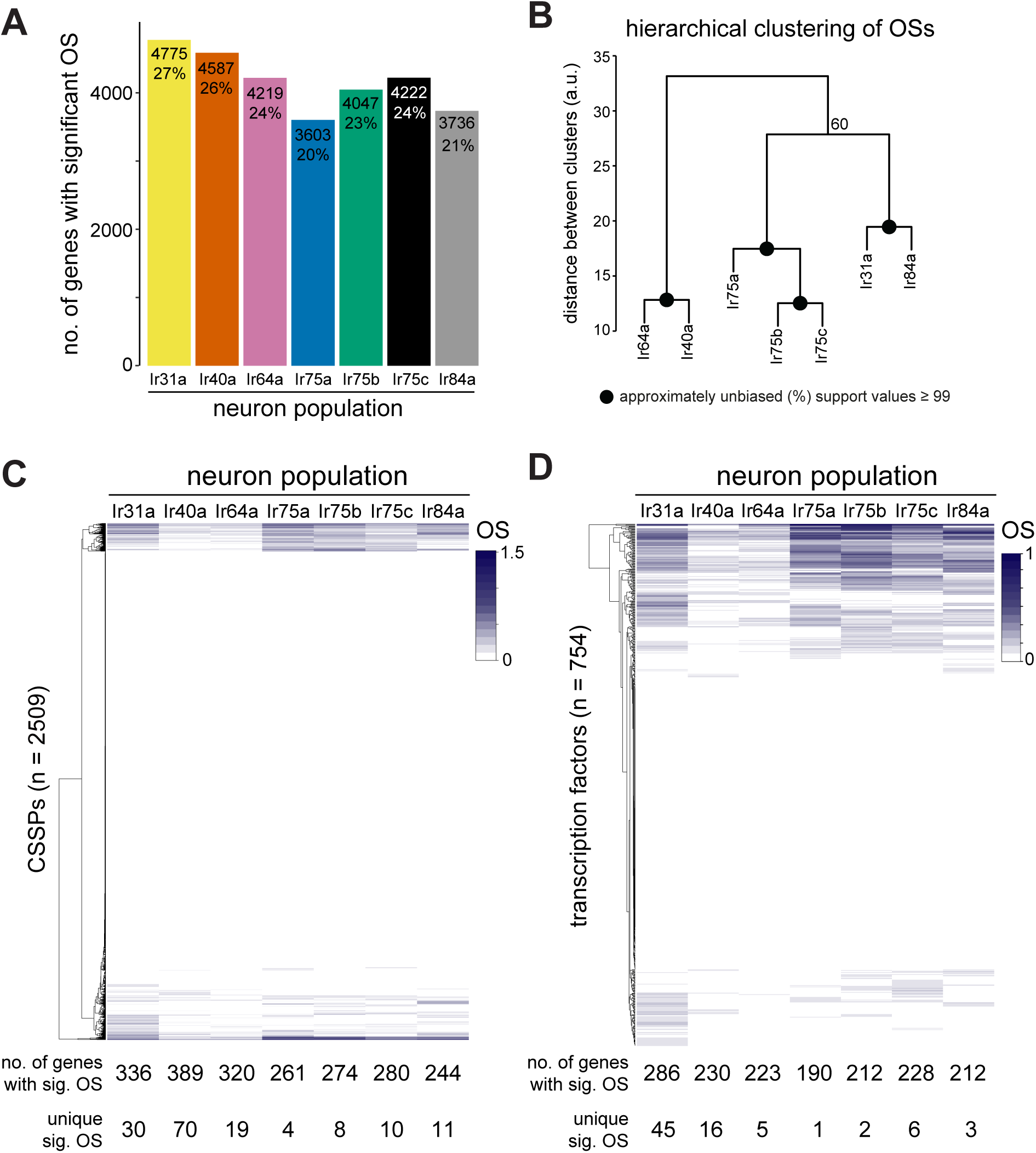
Global analyses of TaDa datasets. (A) Bar plot summarizing the number and percentage of the total genes in the genome that have significantly positive OSs within each Ir neuron population (out of 17,611 genes in the *D. melanogaster* genome). (B) Hierarchical clustering of the seven Ir neuron populations based upon the OSs of all genes. All nodes had strong support except for the one denoted with an Approximately Unbiased *p*-value of 60. (C) Heatmap displaying the hierarchical clustering of OSs for the 2509 genes encoding cell surface/secreted proteins (CSSPs)^44^. For each population, the total number of genes with a significant OS, and the number of those that are unique to a given population, are shown at the bottom (see also Tables S2-4). (D) Heatmap displaying the hierarchical clustering of OSs for the set of 754 putative transcription factor genes^46^. For each population, the total number of genes with a significant OS, and the number of those that are unique to a given population, are shown at the bottom (see also Tables S5-7).

Different OSN populations are characterized both by the sensory receptors they express and by their distinct morphological properties, notably their axonal projections to discrete glomeruli within the antennal lobe. Genetic studies have identified several secreted and transmembrane proteins that play roles in different steps of the neuronal guidance process^3,8,9^, although our understanding remains incomplete and many key molecules likely remain to be discovered. To gain insight into the potential complexity and neuron-type specificity of the cell surface proteome, we examined OSs of ∼2500 cell surface/secreted proteins (CSSPs) annotated in the genome (from the FlyXCDB database^44^) (Figure 2C and Table S2). Each neuronal population is predicted to express 244-389 CSSP genes (Table S3): these are likely to encode proteins involved in diverse structural and functional properties of OSNs, including sensory transduction, dendrite morphogenesis, axon guidance, and synaptic maturation and transmission. We did not attempt to further categorize the expressed genes, however, as most of them have unknown *in vivo* roles. While each neuron population had a unique combination of PolII-occupied CSSP genes, only a small number of these were unique to a single dataset (Figure 2C; Table S4). One exception is the Ir40a population (70 uniquely occupied genes); this observation could reflect the distinct sensory modality mediated by these neurons (hygrosensation) and/or their unusually distributed glomerular innervations in the antennal lobe^22^.

The expression of both chemosensory receptors and guidance molecules depends upon regulatory networks of transcription factors^2,45^. We therefore assessed global patterns of PolII occupancy of the set of 754 predicted *Drosophila* transcription factors across the seven Ir populations (using the FlyTF database^46^; Table S5). Half of these genes (379/754) displayed significant occupancy in at least one population of Ir neuron, with individual OSN classes displaying 190-286 occupied transcription factor genes (Figure 2D and Table S6-S7). There was considerable variation in Dam:PolII occupancy of transcription factor genes across the neuron populations (Figure 2D), consistent with each neuronal subtype possessing a unique and complex gene regulatory network to support its specific differentiation properties.

Many of the transcription factors with known function in OSN development are broadly-expressed across OSNs, such as Acj6, Fer1 and Onecut^45,47^; consistently, these genes display positive OS in all seven neuron populations (Table S6). One exception is Engrailed (En), which is expressed and required in just a subset of OSNs^48,49^. In our TaDa datasets, we observed that *en* displays significant OSs in Ir31a, Ir40a and Ir64a populations (Figure S2A), consistent with the robust detection of En protein in only these three Ir neuron classes (Figure S2B-C).

### Comparative chromatin accessibility in OSNs

We next performed CATaDa analysis through investigation of the Dam-alone datasets, by using these to generate chromatin accessibility maps similar to ATAC-seq and FAIRE-seq^15^. Genome-wide visualization of the peaks of methylated sequences across the genome (see Methods) revealed heterogeneous patterns of chromatin accessibility, with the expected overall decrease in open chromatin towards the heterochromatic centromeric regions of the chromosome arms (Figure 3A).

**Figure 3.**
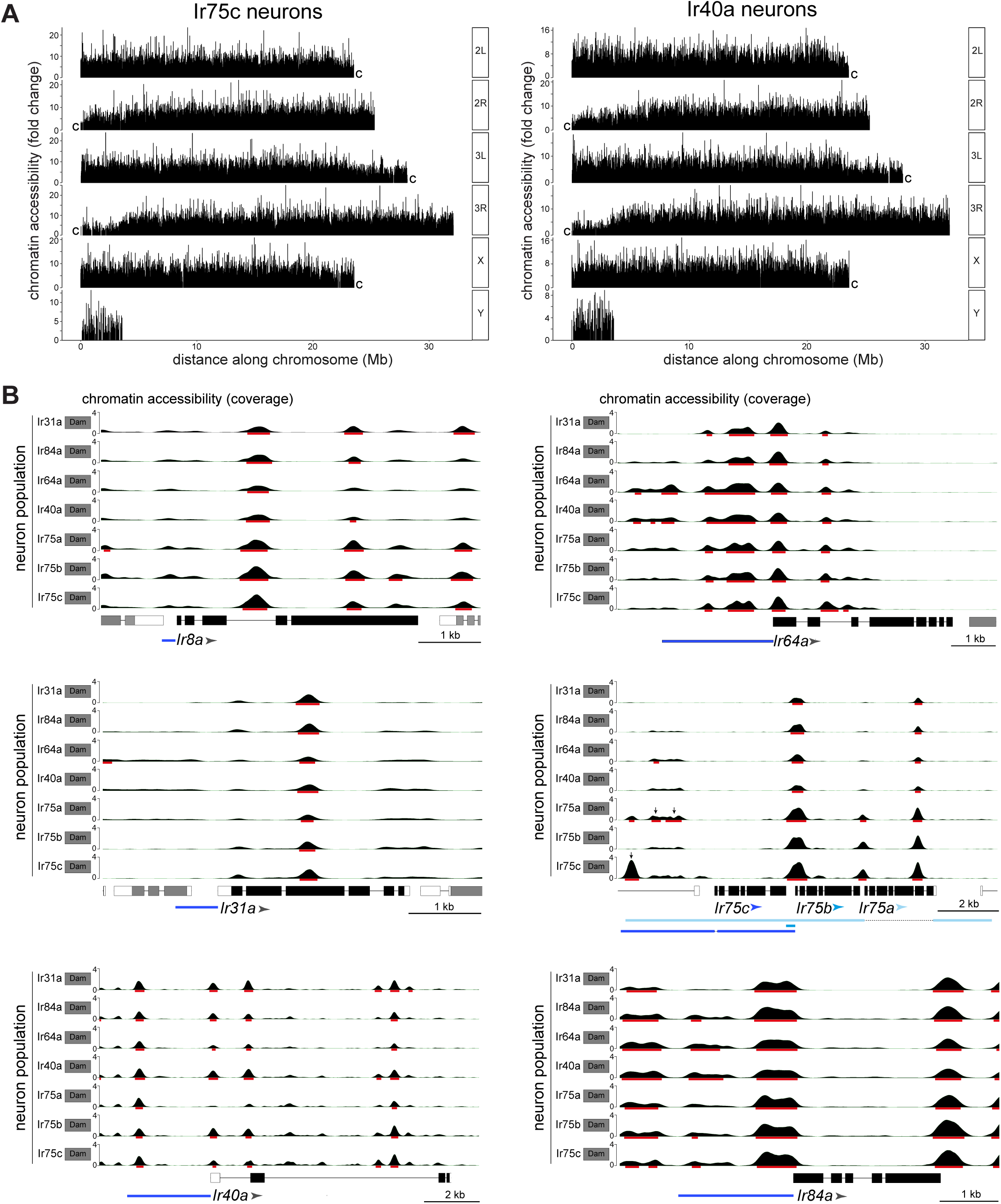
CATaDa reveal global similarity in chromatin architecture at *Ir* genes across Ir neuron populations. (A) Comparison of chromatin accessibility peak locations (q < 0.05) over the *D. melanogaster* genome for two example neuron populations, Ir75c and Ir40a. The y-axis represents the fold enrichment for the peak summit against random Poisson distribution of the small local region (1000 bp). 2L/3L and 2R/3R refer to left and right arms of chromosomes 2/3; “c” at the end of each arm indicates the centromeric (heterochromatic) end of the arm (the Y chromosome is mainly heterochromatic). (B) Plots of *Ir* gene exons are shown in black, and UTRs in white; exons of flanking genes are shown in gray. The arrowheads indicate the direction of *Ir* transcription. The minimal defined regulatory sequence to recapitulate *Ir* expression is indicated with blue bars (color-coordinated with the arrowheads for *Ir75a, Ir75b* and *Ir75c*)^*22,25*^. The y-axis scale shows the CATaDa accessibility at the genomic region with RPGC normalization method. The red bars are the significant peaks with q < 0.05. In the *Ir75a/Ir75b/Ir75c* genomic region, the arrows mark significant peaks that are unique to the corresponding Ir neuron populations, and which are contained with known regulatory regions of these *Ir* genes.

To examine chromatin accessibility patterns at a gene level, we focused on CATaDa signals at the *Ir* loci that distinguish different populations – i.e., the seven tuning *Ir* genes and *Ir8a* – whose cell type-specific transcriptional activity is well-established^25,29,30^. For each gene, peaks were present at the presumed promoter region, as well as in upstream regions, which may represent enhancers. Comparisons across neuron populations revealed that peak distribution was very similar across all eight genes (Figure 3B). This observation indicates that neuron-specific differences in *Ir* gene transcription are not due to differences in promoter accessibility and also not reflecting dramatic differences in enhancer accessibility. The only exceptions were the presence of peaks upstream of *Ir75a* and *Ir75c* that were unique to their respective neuron populations (q-value < 0.05). Notably, these peaks lie within sequences that recapitulate receptor expression patterns in reporter transgenes^20,22,25^ (Figure 3B), raising the possibility that they reflect enhancers that are accessible/functional only in these Ir neuron populations.

### Identifying genes displaying differential expression in Ir populations

The analyses of genes encoding chemosensory receptors, transcription factors and CSSPs demonstrate the use of TaDa OSs to identify genes that are expressed (or not expressed) within specific OSN populations. While differences in OS for a given gene across populations may suggest heterogeneous expression levels, we sought to test for PolII occupancy differences between populations. We implemented a different analysis method that used variation in read-depth in the triplicate paired Dam-alone/PolII:Dam TaDa datasets along the gene body (Figure 4A and Methods). We identified candidates that had at least two GATC motifs for which the excess of PolII:Dam read-depth compared to the Dam-alone read-depth significantly varied in the same direction between two or more population datasets (see Methods). With this criterion, 1694 genes exhibited significant variation across the seven populations, indicating that only a relatively small proportion of genes (∼8%; Dataset S2) differ among these neuron types. This number is likely to be an overestimate as it includes overlapping and/or intronic genes that, as described above, cannot be parsed further using TaDa data. Importantly this subset encompasses all of the expected *Ir* genes (see Figure 4D and Dataset S2). To examine how each individual Ir neuron dataset varied in comparison to the six others, we also quantified significant differences between each pair of neural populations (Figure 4B; Dataset S3). Despite the methodological differences with the TaDa analysis that produces OSs, the proportion of genes contributing to the total number of pairwise difference approximates the clustering of global OSs (Figure 2B). For example, comparisons among the closely clustered Ir75a/Ir75b/Ir75c datasets result in very few of the total differences (0.4-1.7%). Conversely, slightly more than half (51%) of the total differences in the Ir84a dataset arise from comparisons with the distantly clustered Ir40a dataset.

**Figure 4.**
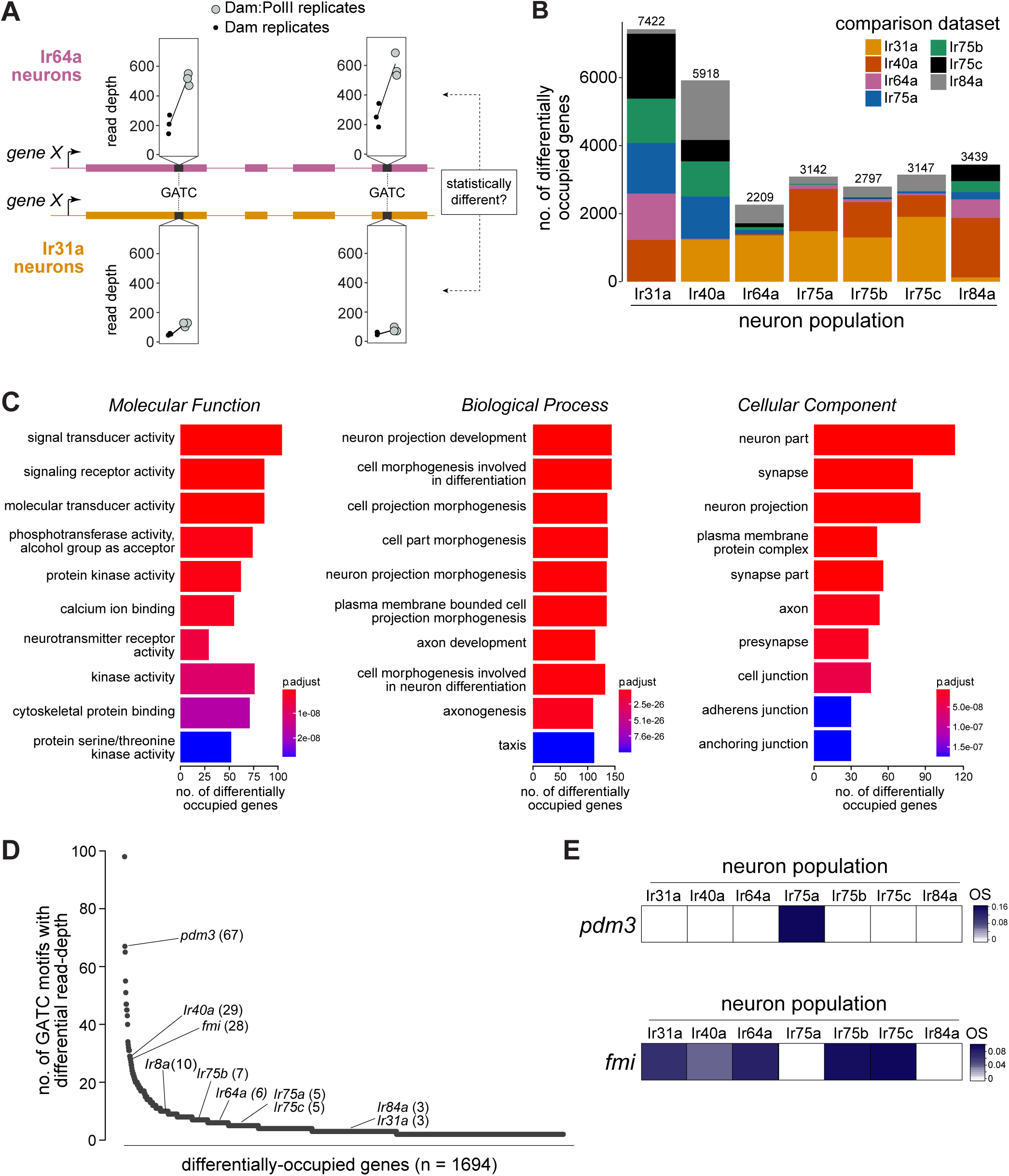
Differentially-occupied genes across Ir populations. (A) Schematic illustrating the identification of a differentially-occupied gene between two Ir populations by using variation in read depth at GATC motifs (see Methods). In this example, occupancy of “*gene X*” is compared between Ir64a neurons and Ir31a neurons. At GATC motifs (black bars), a test is applied to determine if significant variation exists in the relative read depths (in triplicate Dam:PolII versus Dam-alone experiments) between these two neuron populations. Fictitious data for two GATC motifs are depicted: for both, the relative read depth in the Dam:PolII experiments (compared to the Dam-alone experiments) is greater in the Ir64a neurons. (B) Stacked bar plots of the numbers of genes in the genome that are differentially occupied among Ir neuron populations based on pairwise tests (see Methods). The colors indicate proportions of differentially occupied genes that arise from comparisons with the other six neuron populations. The figures above the bars indicate the numbers of genes across all comparisons. (C) Gene Ontology (GO) analysis of the 1694 differentially-occupied genes, showing the top ten over-represented GO terms. The x-axis represents the number of genes annotated to a particular GO term in the input subset. The *p*-value indicated by the colored bar is the probability of observing at least the same number of genes associated to that GO term compared to what would have occurred by chance. (D) Ranking of the 1694 differentially-occupied genes ordered by the number of GATC motifs contributing to their between-neuron differences (uncorrected for gene length). (E) OS heatmaps (calculated as in Figures 1-2) of *pdm3* and *fmi* in the seven Ir neuron populations.

We investigated the general characteristics of the 1694 differentially-occupied genes by performing Gene Ontology analyses, finding highly significant enrichment in neural, neural development, and signaling categories (Figure 4C). This enrichment is consistent with these genes underlying the known unique physiological and anatomical properties of these neuron types, and indicates that our comparative TaDa approach would be valuable in identifying candidates that contribute functionally to these unique properties.

To establish a priority list for follow-up experiments, we ordered these genes by the total number of differentially-occupied GATC motifs, reasoning that this would highlight those displaying differential occupancy along a broad region of the gene body (Figure 4D and Table S8). While this “ranking” is biased towards longer genes (which on average will have more GATC motifs), we emphasize that this does not exclude interest in genes with fewer GATC motifs; for example, *Ir* genes are distributed widely in this list (Figure 4D). We subsequently focused on experimental validation of the top-ranked transcription factor gene, *pou domain motif 3* (*pdm3*), and the top-ranked CSSP gene, *flamingo* (*fmi*; also known as *starry night* (*stan*), flybase.org/reports/FBgn0024836), as described below.

### Pdm3 acts as a genetic switch to distinguish Ir75a and Ir75b neuron fate

The robust signal displayed by *pdm3* for between-population differences is consistent with it having a significant OS only in Ir75a neurons (Figure 4E). Notably, from the pan-transcription factor survey, *pdm3* is the sole gene that is uniquely occupied in this neuron population (Figure 2D and Table S4). Pdm3 is a nervous-system enriched transcription factor^50-52^, although its precise *in vivo* binding specificity is unknown. In another olfactory organ, the maxillary palp, Pdm3 is expressed in multiple classes of Or neurons and functions in controlling receptor expression and axon guidance^52^, but its role in Ir neurons is unknown. Using Pdm3 antibodies, we first analyzed protein expression in antennae in which we co-labeled each of the Ir populations with the driver lines used in the TaDa experiments. Only Ir75a neurons displayed robust nuclear Pdm3 immunofluorescence, although a subset of Ir40a neurons had above-background signals (Figure 5A-B).

**Figure 5.**
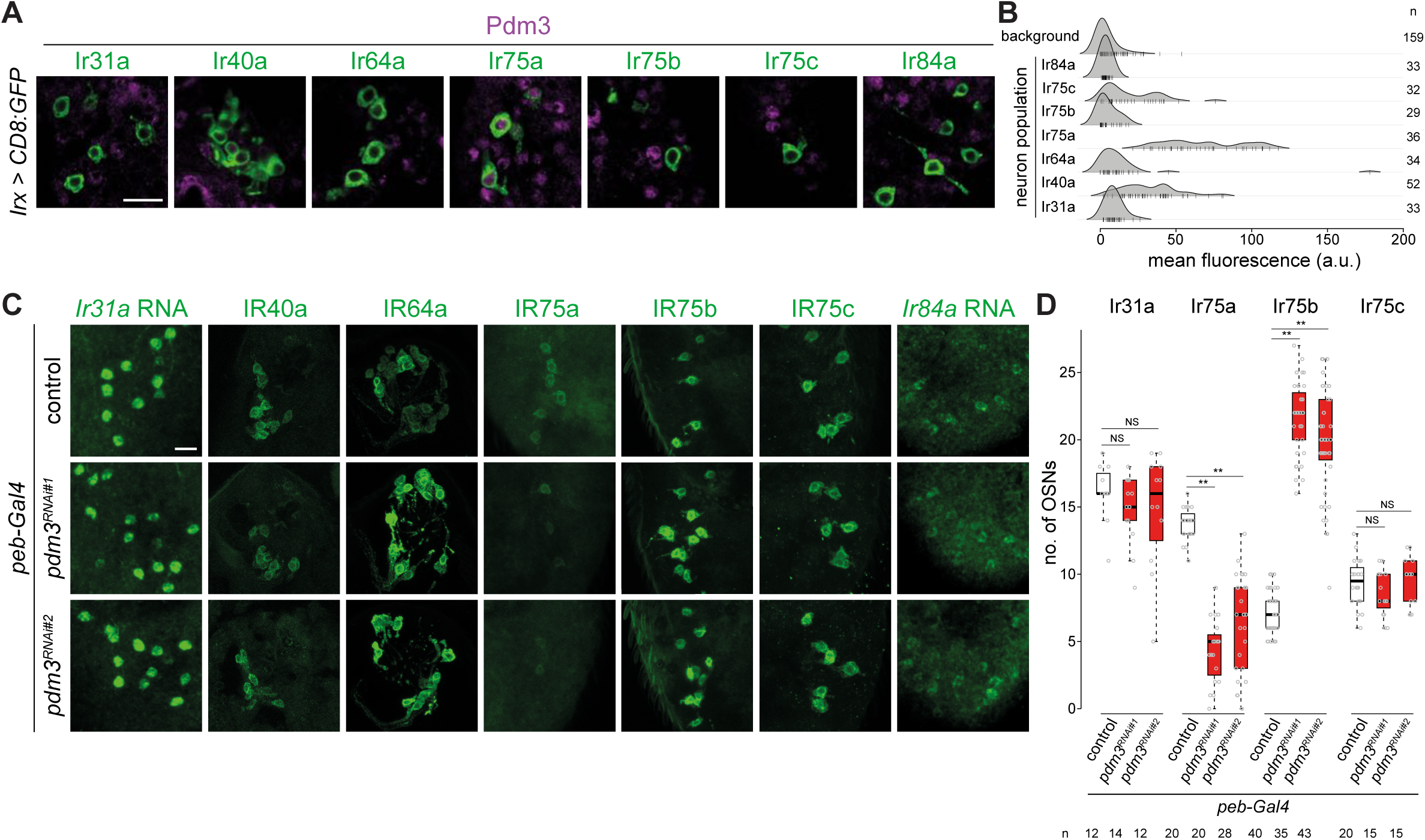
Heterogeneous expression and function of Pdm3 in Ir neurons. (A) Immunofluorescence for Pdm3 and GFP on antennal sections of animals in which the indicated Ir neuron populations are labeled with a CD8:GFP reporter. Genotypes are of the form: *Irxx-Gal4/+;UAS-mCD8:GFP/+* or, for Ir75a and Ir75c neurons, *UAS-mCD8:GFP/+;Irxx-Gal4/+*. Scale bar = 10 µm. (B) Density plots for Pdm3 immunofluorescence signals (arbitrary units, a.u.) quantified from Ir neuron nuclei (each represented by a vertical line) in the genotypes in (A) and from background signals pooled across genotypes (see Methods). Most signals from Ir neuron nuclei fall within the range of background signals; Ir75a neurons have consistently higher levels of immunofluorescence. Sample sizes are shown to the right of the plot. (C) Immunofluorescence or RNA FISH for the indicated IRs on whole-mount antennae (or antennal sections for IR40a and IR64a, due to poor antibody penetration of whole-mount tissue) of control animals (*peb-Gal4,UAS-Dcr-2/+;+/CyO*) or two independent *pdm3* RNAi lines (*peb-Gal4,UAS-Dcr-2/+;UAS-pdm3*^*JF02312*^*/CyO* (RNAi#1) and *peb-Gal4,UAS-Dcr-2/+;UAS-pdm3*^*HMJ21205*^*)/CyO* (RNAi#2)). Scale bar = 10 µm. (D) Quantification of neuron number for the indicated Ir neuron populations for the genotypes shown in (C). In this and other panels, boxplots show the median, first and third quartile of the data, overlaid with individual data points. Comparisons to controls are shown for each neuron type (pairwise Wilcoxon rank-sum two-tailed test and *P* values adjusted for multiple comparisons with Bonferroni method, ** *P* < 0.001; NS *P* > 0.05). Sample sizes are shown below the below. Ir40a and Ir64a OSN population sizes could not be quantified because the sections visualized necessarily contain a variable number of neurons; similarly, we could not confidently count Ir84a OSN number because of the weak signal. Nevertheless, when visualized blindly, the control and RNAi genotypes were not distinguishable for any of these neuron populations (assessing the phenotype in antennae of at least 10 animals from two independent genetic crosses).

To examine the role of *pdm3* in Ir neurons we performed transgenic RNA interference (RNAi) using *peb-Gal4*. This driver begins to be expressed very broadly in OSNs after their terminal divisions (∼16 h APF) and throughout the initiation of olfactory receptor expression (∼40-48 h APF)^13,35^. *pdm3*^*RNAi*^ led to loss of most IR75a expression, with the remaining detectable neurons expressing very low levels of this receptor (Figure 5C-D). By contrast, robust expression of all other Irs was maintained (Figure 5C-D). We noted, however, that the number of cells expressing IR75b increased in *pdm3*^*RNAi*^ antennae (Figure 5D). These observations suggest that Pdm3 functions (directly or indirectly) to promote IR75a expression and repress IR75b expression.

The novel IR75b-expressing cells in *pdm3*^*RNAi*^ antennae are mainly located in the proximal region of the antenna near the sacculus, where Ir75a neurons are normally found (Figure 6A). This distribution suggested that loss of Pdm3 results in a switch of receptor expression from IR75a to IR75b. We investigated this possibility by examining the activity of transcriptional reporters for these receptors, encompassing minimal regulatory DNA sequences fused to *CD4:tdGFP* (hereafter, *GFP*)^25^. Both *Ir75a-GFP* and *Ir75b-GFP* transgenes are faithfully expressed in IR75a and IR75b protein-expressing neurons (Figure 6B-C). In *pdm3*^*RNAi*^ antennae *Ir75a-GFP* expression is lost, while *Ir75b-GFP* is expressed in many ectopic cells (Figure 6B-D).

**Figure 6.**
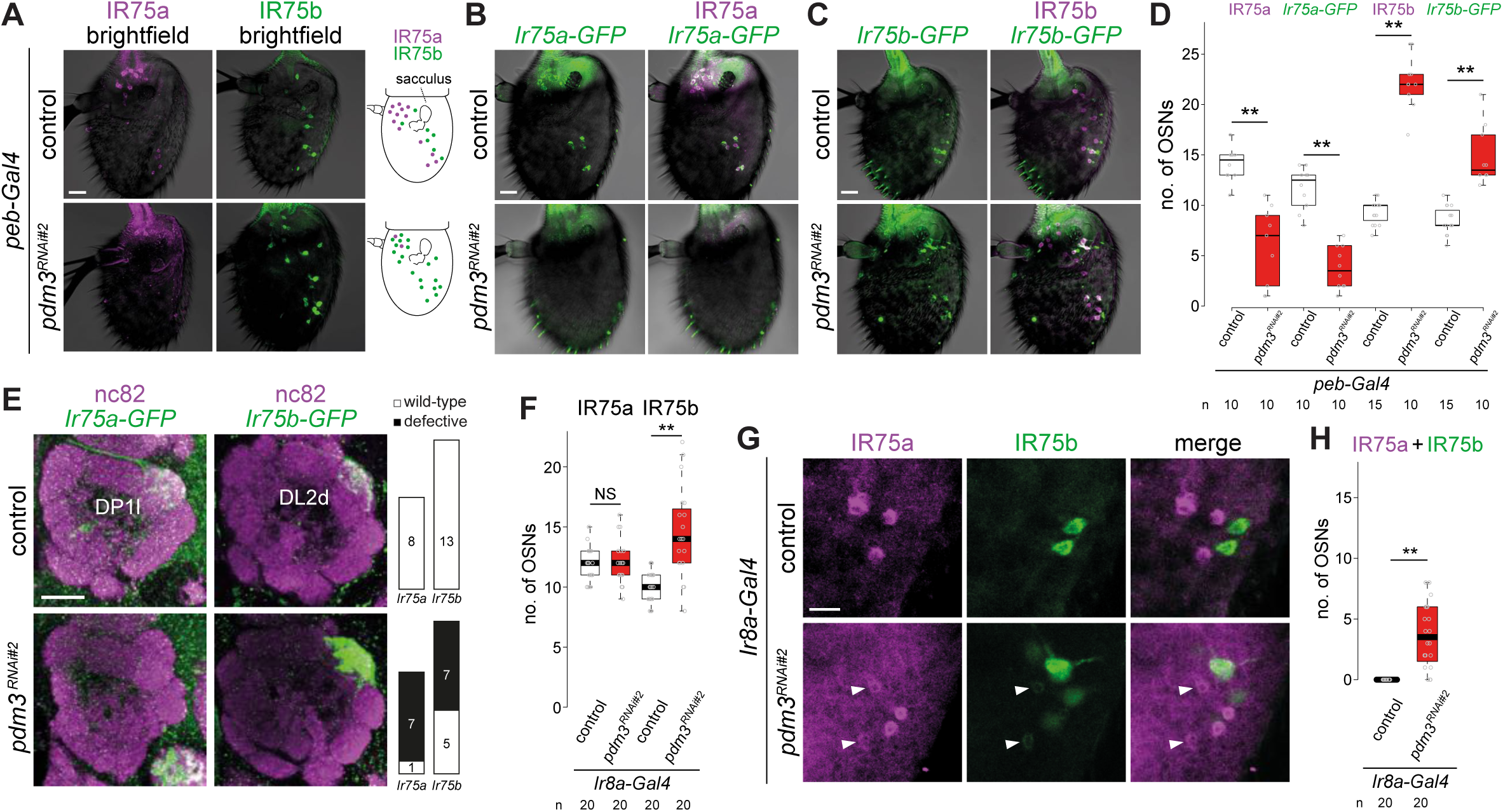
Pdm3 is required to distinguish Ir75a and Ir75b neuron fate. (A) Immunofluorescence for IR75a and IR75b on whole-mount antennae of control (*peb-Gal4,UAS*-*Dcr-2/+*) and *pdm3*^*RNAi#2*^ (*peb-Gal4,UAS*-*Dcr-2/+*;*UAS-pdm3*^*HMJ21205*^*/+*) animals. Scale bar = 10 µm. The schematics on the right summarize the distribution of labeled neurons. (B) Immunofluorescence for GFP and IR75a on whole-mount antennae of control (*peb-Gal4,UAS*-*Dcr-2/+*;*Ir75a-CD4:tdGFP/+*) and *pdm3*^*RNAi#2*^ (*peb-Gal4,UAS*-*Dcr-2/+*;*Ir75a-CD4:tdGFP/UAS-pdm3*^*HMJ21205*^) animals. Scale bar = 10 µm. (C) Immunofluorescence for GFP and IR75b on whole-mount antennae of control (*peb-Gal4,UAS*-*Dcr-2/+*;*Ir75b-CD4:tdGFP/+*) and *pdm3*^*RNAi#2*^ (*peb-Gal4,UAS*-*Dcr2Dcr-2/+*;*Ir75b-CD4:tdGFP/UAS-pdm3*^*HMJ21205*^) animals. Scale bar = 10 µm. (D) Quantifications of the number of neurons that expression *Ir75a*-*GFP* or *Ir75b*-*GFP* in the genotypes shown in (B-C). Comparisons to the controls are shown (pairwise Wilcoxon rank-sum two-tailed test and *P* values adjusted for multiple comparisons with Bonferroni method, ** *P* < 0.001). The increase in number of *Ir75b-GFP* labelled neurons in *pdm3*^*RNAi*^ is lower than the increase in IR75b-expressing neurons, potentially because the transgenic reporter is not fully faithful in this genetic background. (E) Antennal lobe projections of Ir75a and Ir75b neurons in control and in *pdm3*^*RNAi*^ animals. Genotypes are as in (B). Scale bar = 20 µm. Quantifications of projection pattern phenotypes are shown on the right. (F) Quantification of numbers of Ir75a and Ir75b neurons in control (*Ir8a*-*Gal4*/+) and *pdm3*^*RNAi#2*^ (*Ir8a*-*Gal4*/*UAS-pdm3*^*HMJ21205*^) animals. Comparisons to the control are shown for each neuron type (pairwise Wilcoxon rank-sum two-tailed test, ** *P* < 0.001, NS *P* > 0.05). (G) Immunofluorescence for IR75a and IR75b in whole-mount antennae of control and *pdm3*^*RNAi#2*^ animals. Single optical sections are shown, to reveal the weak co-expression of IR75a and IR75b in a subset of cells (arrowheads). Scale bar = 10 µm. (H) Quantifications of number of neurons that co-express IR75a and IR75b. Comparison to the control is shown (pairwise Wilcoxon rank-sum two-tailed test).

We further used these *Ir-GFP* transgenes to examine the glomerular projection of Ir75a and Ir75b neurons in *pdm3*^*RNAi*^ animals. In controls, *Ir75a-GFP* and *Ir75b-GFP* label exclusively the DP1l and DL2d glomeruli, respectively (Figure 6E), concordant with previous analyses^22,25^. In *pdm3*^*RNAi*^ animals the *Ir75a*-GFP DP1l signal is almost completely lost (Figure 6E), consistent with the lack of reporter expression in the antennae. By contrast, the *Ir75b-GFP* signal in the DL2d glomerulus is greatly intensified, and the glomerulus is enlarged (Figure 6E), presumably reflecting the increased number of Ir75b neurons (Figure 6C-D). This simplest interpretation of these observations is that in the absence of Pdm3, Ir75a neurons adopt Ir75b neuron fate encompassing both its receptor identity and glomerular innervation pattern.

While OSN fate is established in early pupae^1-3^, the expression of Pdm3 in Ir75a neurons in adult antennae (Figure 5A-B) suggests that it has a persistent role in these cells. To test this idea, we drove *pdm3*^*RNAi*^ using *Ir8a-Gal4*^29^, which is active only from the second half of pupal development (∼48 h APF) by which time OSN axonal convergence on glomeruli has occurred. In adult *Ir8a>pdm3*^*RNAi*^ antennae, we found that the number of IR75a-expressing cells was unchanged compared to controls, although protein levels were lower in several cells, arguing that Ir75a neuron fate had been largely correctly established (Figure 6F-G). However, as observed with *peb-Gal4*-induced *pdm3*^*RNAi*^, IR75b was still detected in many ectopic cells (Figure 6F). Notably, we detected several cells expressing both IR75b and IR75a, which are never observed in control antennae (Figure 6G-H). These observations indicate that Pdm3 maintains its role in repressing Ir75b expression in mature Ir75a neurons.

### Dynamic, heterogeneous expression of Fmi in OSNs

*fmi* attracted our attention both for its heterogeneous occupancy across Ir neuron populations (Figure 4D-E), and because this gene encodes an atypical cadherin – comprising an extracellular cadherin-like domain and a transmembrane G protein-coupled receptor (GPCR)-like domain – with diverse roles in axonal and dendritic targeting in other regions of the nervous system^53-58^. To characterise the endogenous expression of Fmi in the olfactory system, we used an antibody to profile protein levels in OSNs. We first examined antennae from mid-pupal stages (48 h APF) but observed only trace levels of immunoreactivity across OSN soma (Figure S3A). At 24 h APF this signal was slightly stronger, but much more prominent in the OSN axons as they left the antenna (Figure S3A), suggesting that this protein is predominantly transported to the axonal compartment of these neurons. Indeed, Fmi protein was strongly detected in the OSN axons as they enter the antennal lobe at 24 h APF (Figure 7A). Fmi was still robustly expressed at 48 h APF as these OSNs coalesced onto individual glomeruli with broad, but slightly heterogeneous, expression in glomeruli across the antennal lobe (Figure 7A). Subsequently, Fmi levels began to decrease to undetectable levels in some glomeruli in the adult stage, while reproducibly persisting in others (Figure 7B). Most of these glomeruli are innervated by Or neurons, but we could detect Fmi in DL2d (Ir75b) and DL2v (Ir75c), but not the neighboring DP1l (Ir75a) (Figure 7B), in partial concordance with the TaDa data (Figure 4E).

**Figure 7.**
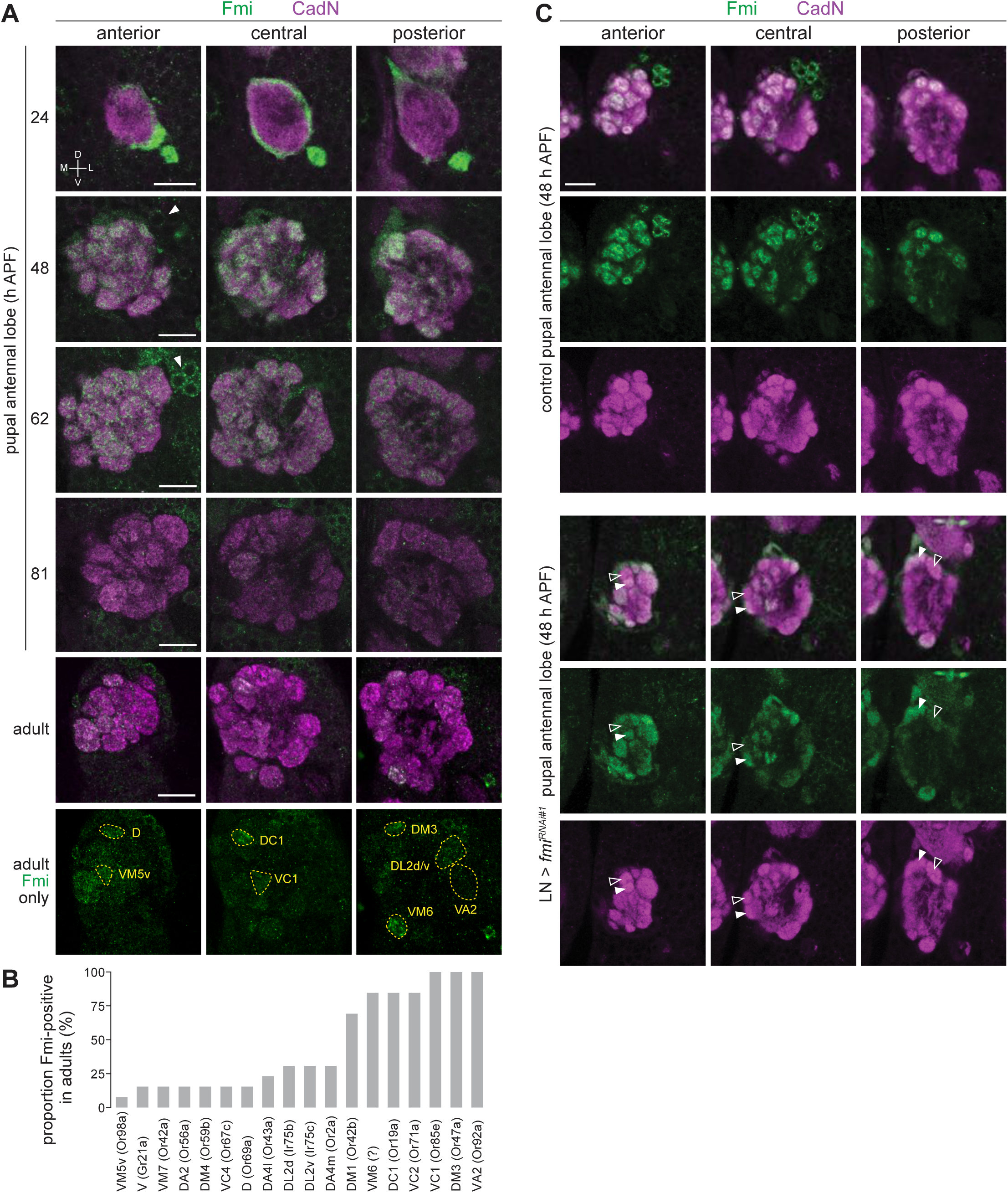
Expression analysis of Fmi in OSNs. (A) Immunofluorescence for Fmi and the ubiquitously-expressed neuronal cadherin (CadN) on whole mount antennal lobes of wild-type (*w*^*1118*^) animals of the indicated age. Three optical sections are shown to reveal the morphology of most glomeruli. Bottom row: Fmi-positive glomeruli were identified in adult antennal lobes based upon stereotyped position and morphology. Dorsal-ventral (D-V) and lateral-medial (L-M) axes are indicated. Scale bar = 20 μm. (B) Histogram of frequency of detectable Fmi immunoreactivity in glomeruli of the adult antennal lobe (n = 14 antennal lobes). (C) Immunofluorescence for Fmi and CadN on control (*1081-Gal4/+;449-Gal4/UAS-Dcr-2*) and *LN>fmi*^*RNAi#1*^ (*1081-Gal4/UAS-fmi*^*KK100512*^;*449-Gal4/UAS-Dcr-2*) animals. Open and filled arrowheads highlight adjacent glomeruli with low and high levels of Fmi staining. Scale bar = 20 μm.

Fmi was also detected in a group of ∼16 cells located on the antero-dorsal region of the antennal lobe from later developmental stages (∼48 h APF) (Figure 7A). These cells are likely to be LNs, based upon their expression of Elav (Figure S3B) and presence of stained processes that innervate the antennal lobe but do not project to higher olfactory centers. Numerous LN subtypes exist, and these often have broad innervation patterns across many glomeruli^59,60^. To examine the specific contribution of OSNs to the Fmi immunoreactivity observed in the antennal lobe, we depleted Fmi in LNs by RNAi using a combination of Gal4 drivers (*1081-Gal4* and *449-Gal4*^60^) that cover the vast majority of these interneurons (Figure S3C). At mid-pupal development (48 h APF), differential glomerular expression of Fmi was much more marked compared to control animals (Figure 7C). Although we could not confidently define all glomerular identities at this earlier developmental stage, prior to full maturation of the antennal lobe, Fmi protein levels did not appear to correspond precisely with the TaDa OSs (e.g., the VL2a glomerulus, innervated by Ir84a neurons, was not devoid of Fmi signal). This observation suggests that Fmi is dynamically expressed in different populations of OSNs during development. We note that the protein observed at 48 h APF must be due to transcription of *fmi* at an earlier time point than captured by our TaDa analysis. Additionally, different post-transcriptional regulation within OSN populations could also contribute to lack of strict correlation between transcript and protein levels with these cells, as noted in other tissues^61^. Nevertheless, this analysis reveals developmentally dynamic and population-specific control of Fmi expression in OSNs.

### Fmi is required for segregation of OSNs into distinct glomeruli

To determine the function of Fmi in the developing olfactory system, we induced *fmi* RNAi in developing antennal tissue first using a constitutive driver (*ey-Flp,act>stop>Gal4*^62^), which should deplete Fmi expression from the earliest stages of development), and examined adult brains stained with the general neuropil marker nc82 (Bruchpilot). The antennal lobes of these animals showed fully penetrant, severe morphological defects: the stereotyped glomerular boundaries observed in the lobes of control animals were largely lacking, with only a few, large glomerulus-like subregions detected (Figure 8A). This phenotype was reproduced with an independent RNAi line that targets a distinct region of the coding sequence (Figure 8A)

**Figure 8.**
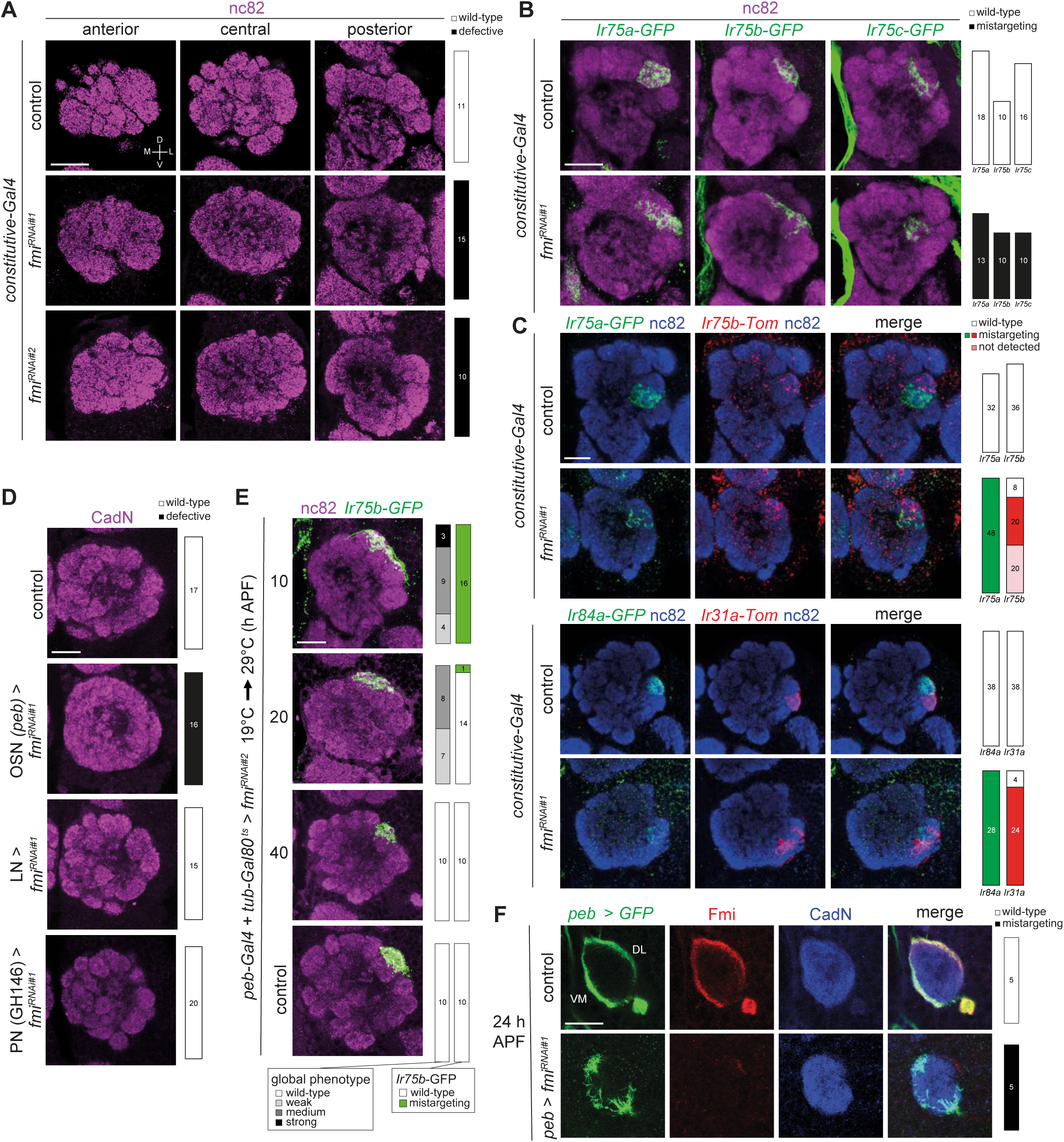
Fmi is required in OSNs for glomerular segregation. (A) Immunofluorescence for nc82 on whole mount antennal lobes of control (*w;ey-Flp/+;act>stop>Gal4/+*), *fmi*^*RNAi#*1^ (*w;ey-Flp/UAS-fmi*^*KK100512*^;*act>stop>Gal4/+*) and *fmi*^*RNAi#*2^ (*w;ey-Flp/+;act>stop>Gal4/UAS-fmi*^*JF02047*^) animals. Three optical sections are shown. Scale bar = 20 μm. Quantifications of wild-type or defective antennal lobe glomerular architectures are shown on the right; the n for each phenotypic category are indicated on the plots. (B) Immunofluorescence for nc82 and GFP on whole mount antennal lobes of control and *fmi*^*RNAi#1*^ animals. Genotypes: *Ir75a*-GFP (*w;ey-Flp,Ir75a-GFP/*[*+* or *UAS-fmi*^*KK100512*^];*act>stop>Gal4/+*), *Ir75b*-GFP (*w;ey-Flp,Ir75b-GFP/*[*+* or *UAS-fmi*^*KK100512*^];*act>stop>Gal4/+*), *Ir75c*-GFP (*w;ey-Flp,Ir75c-GFP/*[*+* or *UAS-fmi*^*KK100512*^];*act>stop>Gal4/+*). Scale bar = 20 μm. (C) Immunofluorescence for nc82, GFP, RFP (detects Tomato (Tom)) on whole mount antennal lobes of control and *fmi*^*RNAi#1*^ animals. Genotypes: *w;Ir75a-GFP,Ir75b-RFP/*[*+* or *UAS-fmi*^*KK100512*^];*ey-FLP,act>stop>Gal4/Ir75b-RFP* (top panels) and *eyFlp,act>stop>Gal4*/*/*[*+* or *UAS-fmi*^*KK100512*^];*Ir31a-RFP,Ir84a-GFP/+* (bottom panels). Scale bar = 20 μm. (D) Immunofluorescence for CadN on whole mount antennal lobes of control (*peb-Gal4*), OSN*>fmi*^*RNAi#1*^ (*peb-Gal4/+;UAS-fmi*^*KK100512*^*/+;UAS-Dcr-2/+*), LN*>fmi*^*RNAi#1*^ (*1081-Gal4/UAS-fmi*^*KK100512*^;*449-Gal4/UAS-Dcr-2)*, PN*>fmi*^*RNAi#1*^ (*GH146-Gal4/+;UAS-fmi*^*KK100512*^*/+*) animals. Scale bar = 20 μm. (E) Immunofluorescence for nc82 and GFP on whole-mount antennal lobes of *peb-Gal4,UAS-Dcr-2/+;Ir75b-GFP/tub-Gal80*^*ts*^;*UAS-fmi*^*JF02047*^*/+* animals in which Gal4-driven RNAi was suppressed until shifting from the permissive to restrictive temperature (19°C > 29°C) for Gal80^ts^ at the indicated time points. Full confocal-stacks are shown. Scale bar = 20 μm. (F) Immunofluorescence for GFP, Fmi and CadN on whole mount pupal antennal lobes (24 h APF) of control (*peb-Gal4/+;;UAS-mCD8::GFP/+*) and OSN*>fmi*^*RNAi#1*^ (*peb-Gal4/+;UAS-fmi*^*KK100512*^*/+;UAS-mCD8::GFP/+*) animals. The merged channels are shown on the right. The dorsolateral (DL) and ventromedial (VM) bundles of OSN axons are indicated. Scale bar = 20 μm.

We further examined this phenotype by visualizing the projection patterns of specific classes of OSNs labeled by receptor promoter-driven fluorescent reporters (Figure 8B-C). Neurons labeled by *Ir75a-GFP, Ir75b-GFP* or *Ir75c-GFP* were no longer constrained to discrete glomeruli (Figure 8B), although the mistargeting was relatively mild within the antennal lobe as a whole. The limited defects of individual neuron populations suggests that the lack of Fmi does not disrupt long-range targeting properties of OSNs, but rather impacts local segregation of OSNs to discrete glomerular targets. Consistently, double labeling of neurons that normally project to adjacent glomeruli (*Ir75a-GFP* and *Ir75b-Tomato* (*Tom*); *Ir84a-GFP* and *Ir31a-Tom*) revealed frequent overlap between these normally-segregated populations (Figure 8C).

The constitutive driver is expressed broadly across the antennal disc^62^. The observed phenotypes could therefore potentially be due to a requirement for Fmi in disc development rather than in OSNs. To limit *fmi* RNAi to OSNs, we used *peb-Gal4*, which produced an equally strong defect in antennal lobe glomerular segregation (Figure 8D). In these animals the LN-expressed Fmi was still detectable but, in contrast to the expression in OSNs, Fmi was apparently homogeneously distributed across antennal lobe glomeruli (Figure S3D). This LN source of Fmi does not appear to contribute to glomerulus formation as *LN>fmi*^*RNAi*^ animals did not exhibit the same antennal lobe morphological defects (Figure 8D). Similarly, when we targeted *fmi* to the synaptic partners of OSNs, the PNs – which pre-pattern the glomerular organization of the antennal lobe^3^ – using the *GH146* driver, antennal lobe morphology appeared normal (Figure 8D). The absence of defects is consistent with the lack of detectable expression of Fmi in these second-order neurons (Figure 7A). Together, these results indicate that Fmi functions principally, if not exclusively, within OSNs to correctly pattern glomerular formation.

To understand the genesis of this defect during OSNs development we used a temperature-sensitive repressor of Gal4 (Gal80^ts^; inactivated at 30°C) to temporally-control the onset of *peb*>*fmi*^*RNAi*^. When animals were shifted to 30°C 10 h APF (prior to initial expression of *peb-Gal4*), all animals displayed antennal lobe morphological defects of varying degrees, and *Ir75b-GFP*-labeled neurons did not coalesce to a discrete glomerular unit (Figure 8E). When shifted at 20 h APF, the severity of the phenotype was diminished (Figure 8E). When shifted at 40 h APF, the antennal lobe and Ir75b neuron targeting were indistinguishable from controls (Figure 8E). These experiments indicate an early developmental function for Fmi.

We examined this early requirement in more detail by imaging the OSN axons as they invade the antennal lobe. During 20-25 h APF, pioneer OSNs reach the antennal lobe in a unique bundle that separates into ventromedial and dorsolateral bundles, which project around the lobe’s border ^9,63^ (Figure 8F). In *fmi*^*RNAi*^ animals, OSNs reach the antennal lobe, but bundle organization is highly disrupted, with subsets of axons apparently defasciculating prematurely in the central area of the lobe (Figure 8F).

Fmi (and its mammalian homologs) have roles in multiple biological processes in the nervous system as well as in the establishment of epithelial planar cell polarity^58,64,65^. Genetic and biochemical studies have identified a number of interaction partners of Fmi in different biological contexts. We tested mutations and/or multiple independent RNAi lines for several of these genes to determine whether they function with this cadherin in OSNs (Figure S4). Mutations in two core components of the planar cell polarity pathway, *frizzled* and *dishevelled*^*66*^, had no apparent impact on antennal lobe formation. Similarly, loss of *frazzled* (encoding a Netrin homolog which function with Fmi in midline axon guidance^67^), *golden goal* (with which Fmi collaborates in photoreceptor axon guidance^68^), or *espinas* (which encodes a LIM domain protein that binds Fmi intracellularly and functions in dendrite self-avoidance^57^) did not reproduce the *fmi* phenotype. Finally, although the intracellular signaling mechanism of Fmi is not well understood, the Fmi GPCR-domain has been suggested to act through the Gαq subunit to regulate dendrite growth^69^; however, we found no evidence that this signaling mechanism operates in OSNs (Figure S4). These observations suggest that Fmi functions through other cellular mechanisms to control OSN axon segregation.

## Discussion

Determination of the molecular composition of neurons is essential to understand the development and function of the nervous system. While scRNA sequencing is an undoubtedly powerful approach to address this challenge^70-74^, it requires often technically-difficult cell isolation, and can be biased towards abundant cell types and highly-expressed genes^75^. Moreover, it is unclear to what extent neurons change their expression properties during tissue dissection (as reported for other types of cell^76^) or through inevitable loss of dendritic and axonal processes (where certain transcripts may be enriched^77^). Our study was initially motivated by the desire to test the TaDa method for directed, genome-wide molecular profiling of very small populations of neurons that are tightly embedded within a highly heterogeneous tissue. Moreover, by applying TaDa to a set of functionally-related OSNs, we aimed to identify cell type-specific differences that would point towards mechanisms underlying the development and evolution of novel neuronal populations.

The unique olfactory receptor expression pattern of OSNs (together with other chemosensory genes) offered several positive- and negative-control loci that allowed us to validate the specificity and sensitivity of TaDa. One caveat emerging from this analysis is that TaDa cannot easily discriminate PolII occupancy of interleaved genes, an issue that may be most pronounced in the relatively gene-dense genome of *Drosophila*. Nevertheless, our results indicate that this method could be effective in defining candidate receptors in neuronal classes for which a genetic driver is available. This possibility is of particular interest in the *Drosophila* gustatory system where many populations of neurons can be labeled and physiologically profiled^28^, but incomplete knowledge of the combinatorial receptor expression properties that are characteristic of this sensory system hampers identification of the cognate sensory receptor(s)^27,78,79^.

Recent work described an scRNA-seq analysis of antennal OSNs at a mid-pupal stage (42-48 h APF)^13^, leading to the identification of transcriptional clusters for 20 molecularly-defined classes of OSNs (a further 13 clusters were not identified). Detailed comparison of these scRNA-seq data and our TaDa datasets is currently difficult, as only two of the scRNA-seq clusters correspond to populations we analyzed by TaDa (Ir75a and Ir84a neurons). Moreover, while the scRNA-seq captures the transcriptional profile during a narrow developmental window, our TaDa experiments integrated PolII occupancy patterns at an OSN population level over several days from the initiation of *Ir-Gal4* expression. Perhaps reflecting this difference in transcriptomic profiling, scRNA-seq detected on average ∼1500 expressed genes (including ∼150 encoding transcription factors) per OSN type, which is lower than the number of occupied genes detected by TaDa (on average ∼4000 per Ir population, including ∼225 transcription factor genes). However, many other experimental and/or analytical differences in the two methods are likely to impact these global numbers.

Genome-wide analysis of TaDa datasets from different Ir populations revealed an intriguing positive relationship between the similarity of global patterns of PolII occupancy between neuronal populations and the phylogenetic distance of the olfactory receptors they express, notably the tandem array of *Ir75a, Ir75b* and *Ir75c* genes. Similar analysis of Or OSN populations is limited by the broad phylogenetic distribution of the ORs whose corresponding neuronal transcriptomes have been identified^13^. However, through examination of these data^13^ we identified two pairs of closely-related (though not tandemly-arranged) *Or* genes whose neuronal transcriptional profiles also cluster (*Or42b*/*Or59b* and *Or9a*/*Or47a*). The simplest interpretation of these observations is that neuronal populations expressing distinct, relatively-recent receptor gene duplicates derive from a common ancestral neuronal population (which may have originally co-expressed the duplicated receptors); these extant populations therefore retain correspondingly close proximity in their transcriptomic profile. The principle of OSN “sub-functionalization” – which has not, to our knowledge, previously been recognized empirically – has important implications for understanding the evolution of novel olfactory pathways^80^.

One important advantage of the TaDa profiling method over scRNA-seq is that it can also provide information on chromatin state, which remains difficult to characterize through standard chromatin immunoprecipitation-based methods^81^. We recognize that CATaDa provides only a relatively coarse-grained view of chromatin state (similar to ATAC-seq^15^), and variant approaches of TaDa (e.g., Dam fusions to chromatin-binding proteins^82^) are very likely to reveal finer-scale differences between populations. Nevertheless, our findings of similar chromatin accessibility at *Ir* genes across Ir populations are in-line with studies in the developing *Drosophila* embryo where chromatin accessibility is comparable between anterior and posterior region, despite substantial differences in patterns of gene transcription across this body axis^83^. We suggest that the specific developmental properties of OSNs are more likely to be driven by the unique combination of TFs they express. Indeed many of the differentially occupied genes between Ir populations encode known or predicted TFs. In agreement, we validate the selective expression and function of Pdm3 in Ir75a neurons, revealing it to be a key factor – and potentially the sole transcription factor – that distinguishes the closely-related Ir75a and Ir75b neuron populations. Future identification of Pdm3 targets in these neurons will be necessary to determine whether it regulates these *Ir*s directly, and to identify other downstream genes that control the innervation of the corresponding neurons to distinct glomeruli.

Our TaDa datasets also revealed a rich diversity in the predicted cell surface proteome of Ir neurons. Because of the temporal breadth of TaDa profiling, these proteins are likely to contribute to differences of these neurons in cilia/dendritic morphogenesis, axon targeting and synapse formation/plasticity. Our initial characterization of one of these, the atypical cadherin Fmi, reveal it to be a dynamically and heterogeneously-expressed, axon-localized protein. Importantly, Fmi is not restricted to Ir neurons, but found throughout the peripheral olfactory system, concordant with the dramatic loss of glomerular architecture in the antennal lobe in the absence of this protein. We have delimited the role of Fmi to initial sorting of OSNs within the antennal lobe; unfortunately, this early requirement makes it difficult to assess the significance of the heterogeneous expression of this cadherin as levels of Fmi cannot be easily discerned in specific OSN populations within the bundle. We have attempted to overexpress Fmi to assess the consequence of this alternative perturbation of Fmi levels on OSN axonal projections but this did not yield overt defects (data not shown). This negative result may simply reflect technical limitation in achieving adequate high-level Fmi overexpression, as reported in studies of this protein in other systems^55^. Differential expression of Fmi has been proposed to be important for its role in photoreceptor axon guidance^54^. As Fmi is thought to function as a homophilic adhesion molecule^66^, we speculate that heterogeneous Fmi-dependent adhesion between OSN populations enables them to sort themselves within the axon bundle and/or defasciculate at specific places, before local, instructive guidance cues direct them to particular locations within the lobe. An adhesion-dependent role for Fmi is consistent with the *fmi* loss-of-function phenotype most closely-resembling that of mutations in *CadN*^*84*^. CadN is however homogeneously expressed across the antennal lobe, and also present in their PN synaptic partners^84,85^. Future work will be required to uncover the special role of the atypical cadherin Fmi in correct segregation of OSNs axons into distinct glomerular subunits.

Beyond further functional characterization of differentially-occupied genes identified in this study, several future applications of TaDa (and CATaDa) in the olfactory system, and other sensory systems, can be envisaged. For example, temporal analysis of PolII occupancy could be refined by restricting Gal4 activity (with Gal80^ts^) to a particular developmental time-window. Mining of our TaDa datasets for population-specific genes may help to design novel OSN lineage-specific driver lines that are expressed earlier in development (of which few are currently known^62^) to profile gene expression patterns prior to onset of receptor expression. Finally, in adult flies, comparison of PolII occupancy patterns in a specific neuron population before, during and after odor exposure could provide insight into neural activity-regulated gene expression and its relevance for sensory adaptation and plasticity.

## Acknowledgements

We thank Cheng-Ting Chien, Ya-Hui Chou, Fisun Hamaratoglu, Vladimir Katanaev, Tadashi Uemura, Liqun Luo, the Bloomington *Drosophila* Stock Center (NIH P40OD018537), the Vienna *Drosophila* Resource Center, and the Developmental Studies Hybridoma Bank (NICHD of the NIH, University of Iowa) for reagents. Sequencing was performed at the Lausanne Genomic Technologies Facility, and the Swiss Institute of Bioinformatics Vital-IT group provided computational resource support. We are grateful to Andrea Brand’s lab for assistance in establishing the antennal TaDa protocol, Tony Southall and Owen Marshall for advice on the DamID analysis pipeline, Simon Anders for suggestions on DESeq2 usage, and members of the Benton laboratory for comments on the manuscript. J.R.A. was supported by a post-doctoral fellowship from the Novartis Foundation for medical-biological Research (12A14), and is currently supported by a Swiss National Science Foundation Assistant Professorship (PP00P3 176956). Research in R.B.’s laboratory was supported by the University of Lausanne, an ERC Consolidator and Advanced Grants (615094 and 833548, respectively) and the Swiss National Science Foundation.

## Author contributions

All authors contributed to experimental design, data analysis and interpretation of data. Figure contributions were as follows: J.R.A. (Figures 1F-G, 2A-D, 4B-E, 5B, 7C, 8E, S1, S2A,C); L.A. (Figures 1B-C, 5A-C, 7C, 8A-E, S2B-C, S3B-C, S4); J.A. (Figures 7A-B, 8A-C, 8E-F, S3D); K.M. (Figure 5C-D, 6, S3A); P.C.C. (Figure 3, 4C). R.B. wrote the paper with contributions and feedback from all other authors.

## Materials and Methods

### *Drosophila* culture

Flies were maintained on a standard wheat flour/yeast/fruit juice diet at 25°C in 12 h light:12 h dark conditions. Published mutant and transgenic *D. melanogaster* strain are described in Table S9. For the temporal control of *fmi* RNAi with Gal80^ts^, flies of the desired genotype were cultivated at 19°C; animals were staged by selecting white pupae (designated as 0 h after puparium formation (APF)) and shifting these animals to 29°C after 10, 20 or 40 h, to permit induction of RNAi at different time-points during pupal development. Unless otherwise indicated for specific experiments, antennae or brains were collected from both sexes.

### Transgenic flies

New transgenes were constructed using standard molecular biological procedures and transgenesis was performed by BestGene Inc. using the phiC31 site-specific integration system. *Ir75a-CD4:tdGFP* (*Ir75a-GFP*) was generated by subcloning the 5’ (7833 bp) and 3’ (1921 bp) genomic sequences from the *Ir75a-Gal4* transgene^22^ to flank *CD4:tdGFP* in *pDESTHemmarG* (Addgene 31221)^86^, and integrated into the attP40 landing site. *Ir75c-CD4:tdGFP* (*Ir75a-GFP*) was generated by subcloning the 5’ (3072 bp) and 3’ (2545 bp) genomic sequences from the *Ir75c-Gal4* transgene^25^ to flank *CD4:tdGFP* in *pDESTHemmarG* and integrated into attP40. *Ir75b-CD4:tdTom* (*Ir75b-Tom*) was generated by cloning the 299 bp genomic sequences immediately upstream of the *Ir75b* start codon (as in the previously-described *Ir75b-GFP* transgene^25^) into *pDESTHemmarR* (Addgene 31222) and integrated independently into attP5 and attP2 (on chromosome II and III, respectively). *Ir31a-CD4:tdTom* (*Ir31a-Tom*) was generated by cloning the 2000 bp genomic sequences immediately upstream of the *Ir31a* start codon, via Gateway recombination, into pDESTHemmarR, and integrated into attP2. *Ir84a-CD4:tdGFP* (*Ir84a-GFP*) was generated by cloning the 1964 bp genomic sequences immediately upstream of the *Ir84a* start codon, via Gateway recombination, into *pDESTHemmarG* and integrated into VK00027.

### TaDa sample preparation

Antennae from ∼2000 flies (∼1-8 days old) of the desired genotype were harvested by snap-freezing flies in a mini-sieve (Scienceware, Bel-Art Products) with liquid nitrogen, tapping flies to detach and collect appendages in a Petri dish under the sieve containing Triton X-100 (0.1%). Third antennal segments were selected under a binocular microscope by pipetting, and transferred to a 1.5 ml Eppendorf tube. After brief rinsing in PBS to remove detergent, antennae (occupying a volume of ∼50-70 μl) were resuspended in 50 µl PBS and homogenized manually with a blue pestle (Sigma Aldrich Z359947-100EA) for at least 2 min. Antennal fragments were rinsed off the pestle with 100 µl PBS and 40 μl EDTA (100 mM) and 20 μl RNase (12.5 μg/μl) (Qiagen 19101) were added to the antennal lysate, mixed and allowed to sit for 3-5 min. 20 μl Proteinase K (Qiagen DNeasy Blood & Tissue Kit 69504) and 200 µl Buffer AL were added, and mixed by gently pipetting up and down with a blue tip ∼50 times, before incubation in a 56°C heat block for 30 min. The digested lysate was centrifuged at 10,000 rpm for 30 s and the supernatant transferred to a new tube and allowed to cool before proceeding with DNA extraction.

Subsequent sample processing steps for TaDa – i.e., *DpnI* treatment (which cuts at adenine-methylated GATC sites), DamID adaptor ligation, *DpnII* treatment (which cuts at non-methylated GATC sites, thereby digesting unmethylated DNA fragments), and PCR amplification and purification of methylated GATC fragments – were performed essentially as described^87^. PCR products were sonicated using a Covaris S220 to obtain ∼300 bp average fragment size; DNA size and quality were verified by Qubit quantification. DamID adaptors were removed by *Sau3AI* digestion. Truseq Nano libraries were prepared at the Lausanne Genomic Technologies Facility and sequenced (SR 100 bp) on a HiSeq 2500.

### TaDa data analysis

To test for an enrichment of the genome occupancy of Dam:PolII relative to Dam alone, we used the “damidseq pipeline” v1.4^88^, with release 6 of the *D. melanogaster* genome as the reference. Within this pipeline, Bowtie v2.2.6^89^ was implemented to align the Illumina sequence reads to the reference, and data processing called on SAMtools v1.2^90^ and BEDTools v2^91^. This pipeline outputs log2 ratio files in bedgraph format. We calculated Pearson’s Correlation Coefficient among triplicate datasets based on the bedgraph files, and also used them to generate a single averaged occupancy scores file for each of the seven neuron data sets. Reproducibility among all seven triplicate datasets (average *r*^*2*^ = 0.85) was consistent with previous TaDa studies^14^. The files containing the averaged OSs were then inputted into the “polii.gene.call” script (https://github.com/owenjm/polii.gene.call) for estimating an FDR for each gene’s occupancy score. A file detailing the workflow and corresponding code is available on GitLab (https://gitlab.com/roman.arguello/ir-tada).

The set of genes encoding cell surface and secreted proteins (CSSPs) were based on a custom filtered set from the *D. melanogaster* extracellular domain database (FlyXCDB, vFB2015_03^44^) (Table S2). The set of genes encoding transcription factors used in the study were based on a custom filtered set from the FlyTF.org database v2^46^ (Table S5).

For testing the hypothesis that OSs are equal among datasets (neuron populations), we used DESeq2^92^. Unlike the DamID pipeline, this approach focused on individual GATC fragments within each gene. To quantify read coverage of each fragment, we used the same bam files that were outputted by the DamID pipeline (above). We converted these to bed files using BEDTools v2 “bamToBed” utility and calculated the coverage of GATC fragments using the Bedtools’ “coverage” utility (Dataset S4). Differential occupancy was tested over the full dataset using a Likelihood Ratio Test, where the full model specified neuron populations and DamID condition (Dam:PolII vs. Dam-alone) as factors, and an interaction term (between DamID condition and cell population): ∼ cell.pop + cond + cond:cell.pop. The reduced model removed the interaction term: ∼ cell.pop + cond. We also carried out pairwise tests using a Wald Test within DESeq2. Genes that were identified as candidates for differential expression were required to have ≥2 GATC fragments contributing to the difference (that were also significantly PolII occupied).

Gene Ontology (GO) enrichment analyses were performed on the subset of genes that showed differential occupancy using clusterProfiler (v3.10.1)^93^. We compared these genes against the three independent, controlled vocabularies provided by GO Consortium that model Biological Processes, Molecular Function and Cellular Component^94^.

### CATaDa analysis

CATaDa was performed on the seven populations of neurons using the same sets of sequencing reads (i.e., Dam-alone data) acquired for TaDa analysis. The bioinformatics pipeline followed the general concepts of a previously described pipeline^15^, but with modifications. For each neuron population, the genomic sequencing reads from all three biological replicates were aligned to release 6.21 of *D. melanogaster* genome with “very-sensitive” settings (Bowtie2 v2.3.0). Following pooling of aligned reads from the replicates (samtools v1.7), the chromatin accessibilities were represented as the coverage of reads per bin of 10 bases, scaled to 1× average genomic coverage (∼142.57 Mb) and averaged over a sliding window of 100 bases (bamCoverage, version 3.1.3). The chromatin accessibility profile was plotted on genome browser tracks with pyGenomeTracks v2.1. Significant accessible regions were identified with peak-caller (MACS2, v2.1.1.20160309) at a FDR cutoff (q-value) of 0.05. The correlation clustering of peaks among all assayed neuron populations were analyzed with multiBamSummary v3.1.3 on Pearson method and bin size of 3000 bases.

### Histology

Immunofluorescence and RNA FISH on whole-mount antennae or antennal cryosections were performed as described^95^. Immunofluorescence on whole-mount brains was performed essentially as described^22,96^. Antibodies used are listed in Table S10. RNA probes for *Ir31a* and *Ir84a* were previously described^30^.

Quantification of En and Pdm3 immunofluorescence signals in OSN soma nuclei (Figure S2B-C and Figure 5A-B) was performed in Fiji^97^. In brief, regions-of-interest (ROIs) were manually defined using the GFP signal (which circumscribes the nucleus) in a single optical slice where the cross-section of a given nucleus has the largest area, followed by automated measurement of the average En or Pdm3 immunofluorescence signal within this ROI. Background fluorescence signals in the tissue were measured in the same way within equivalently-sized, arbitrarily-chosen ROIs that did not overlap with GFP signals or any other OSN nuclei (visualized with DAPI).

### Statistics statement

Unless stated otherwise, statistical significance thresholds were *p* < 0.05 and the tests carried were two-tailed. Measurements were taken from distinct samples, unless otherwise stated, along with a corresponding correction for multiple tests

## Data availability

TaDa sequencing data and experimental information have been deposited in the ArrayExpress database at EMBL-EBI (www.ebi.ac.uk/arrayexpress) and will be available under accession number E-MTAB-8935. Raw image data for Figures 1, 5-8, S2-S4 are available upon request.

## Code availability

A detailed workflow for the TaDa analyses and associated code are available at GitLab (https://gitlab.com/roman.arguello/ir-tada).

## Biological material availability

All unique biological material generated in this work is available from the corresponding author upon request.

## Supplementary Information

### Supplementary Figure Legends

**Figure S1.**
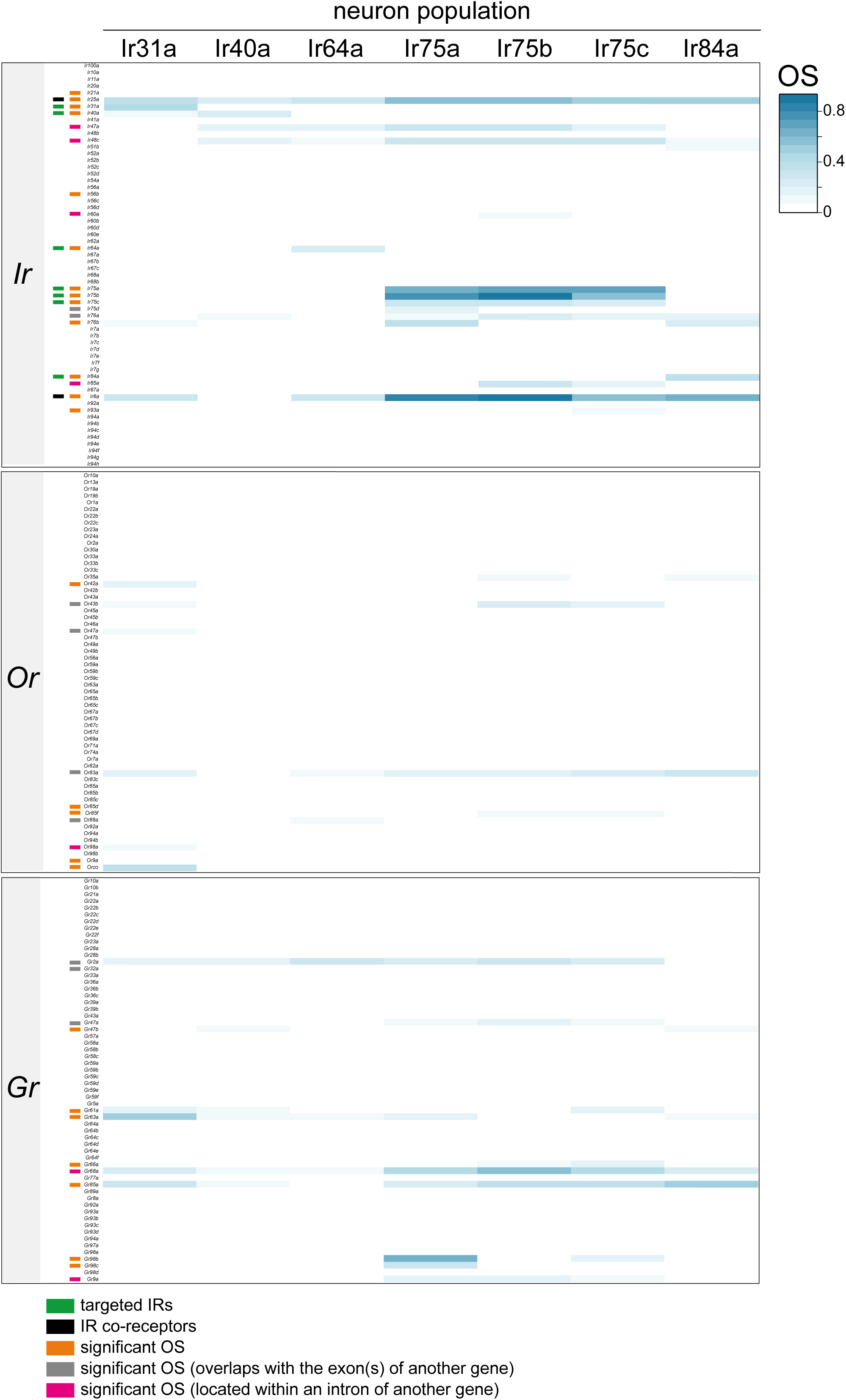
Chemosensory receptor gene Occupancy Scores across Ir populations. Heatmap of OSs for members of *Ir, Or* and *Gr* gene families across the seven Ir neuron populations. Information on individual genes displaying significant OSs in one or more populations is shown with colored bars to the left (key is at the bottom). See also Table S1.

**Figure S2.**
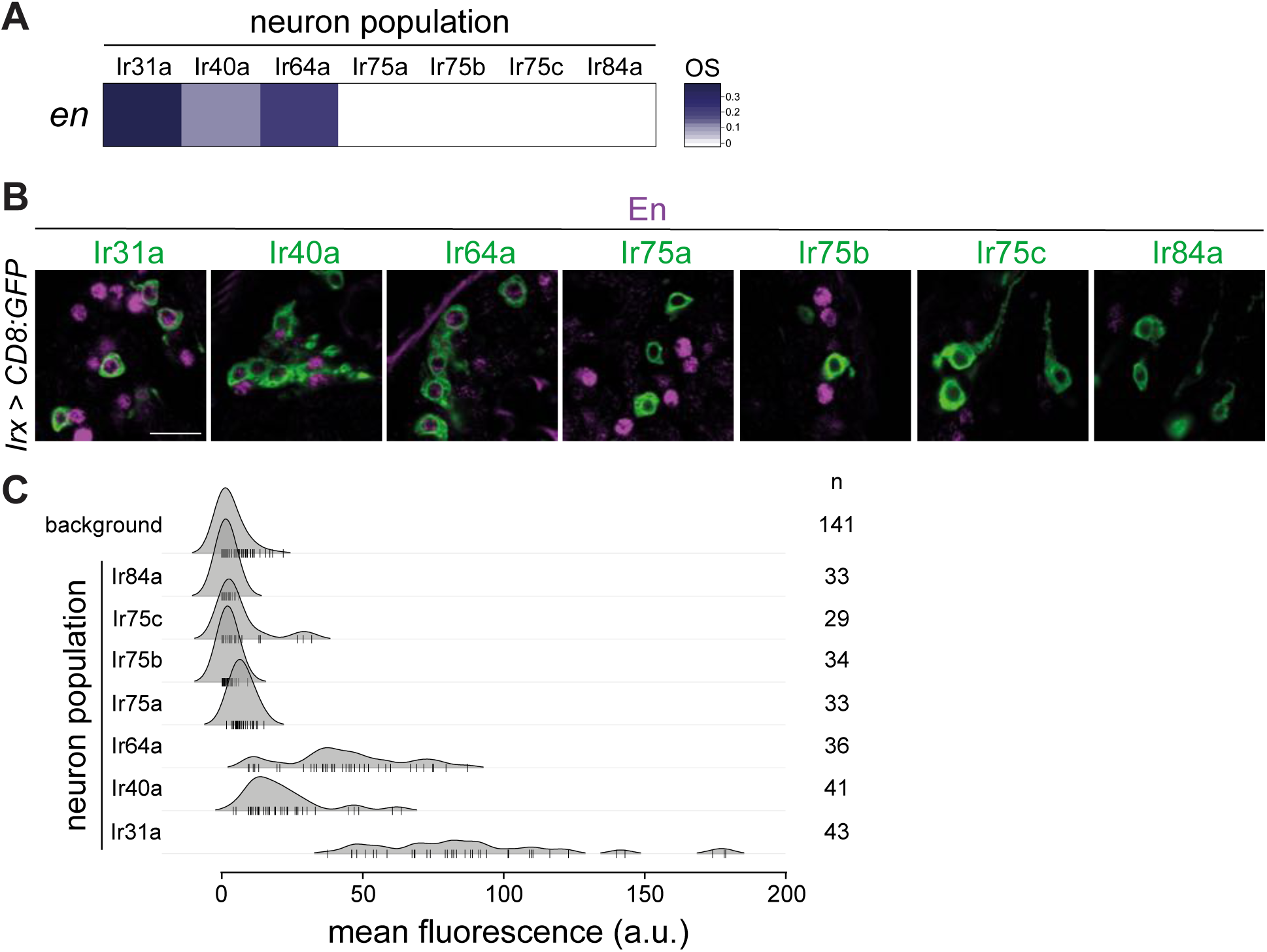
Heterogeneous expression of Engrailed in Ir neuron populations. (A) OS heatmap of *en* in the seven Ir neuron populations. (B) Immunofluorescence for En (magenta) and GFP (green) on antennal sections of animals in which the indicated Ir neuron populations are labeled with a CD8:GFP reporter. Genotypes are of the form: *Irxx-Gal4/+;UAS-mCD8:GFP/+* or, for Ir75a and Ir75c neurons, *UAS-mCD8:GFP/+;Irxx-Gal4/+*. Scale bar = 10 µm. (C) Density plots for En immunofluorescence signals (arbitrary units, a.u.) quantified from Ir neuron nuclei (as in Figure 5B). Most signals from Ir75a, Ir75b, Ir75c and Ir84a neuron nuclei fall within the range of background signals; Ir31a, Ir40a and Ir64a neuron nuclei display shifted distributions.

**Figure S3.**
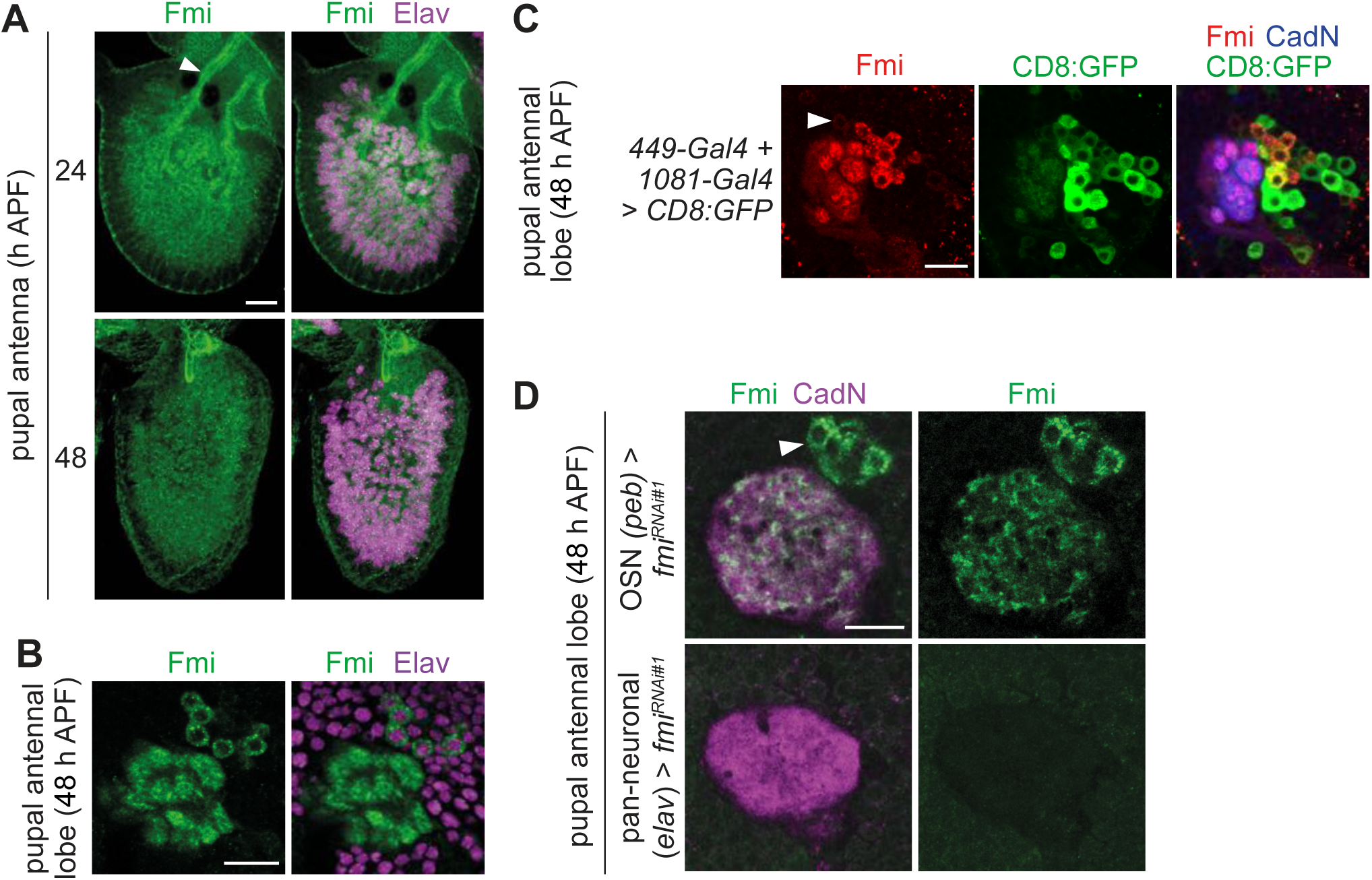
Characterization of Fmi expression in antennae and local interneurons. (A) Immunofluorescence for Fmi and Elav on whole-mount antennae of control animals (*w;Ir8a-Gal4,UAS-RedStinger/CyO*; note the RedStinger reporter is not shown in these images) at 24 and 48 h APF. Partial optical stacks are shown. The arrowhead points to the antennal nerve. Scale bar = 20 μm. (B) Immunofluorescence for Fmi and Elav on whole-mount antennal lobes of 48 h APF control (*w*^*1118*^) animals. A partial optical stack is shown, revealing expression of Elav in the Fmi-positive soma adjacent to the antennal lobe. Scale bar = 20 μm. (C) Immunofluorescence for Fmi, GFP and CadN on a whole-mount antennal lobe of a 48 h APF animal in which Fmi-expressing LNs are labeled with the combined *49-Gal4* and *1081-Gal4* drivers (*1081-Gal4/+;449-Gal4/UAS-mCD8:GFP*), which were identified by screening a panel of LN drivers^1^. A partial optical stack is shown. We distinguish two types of Fmi-positive neurons: 100% of strongly-expressing neurons (n = 194, 7 brains) are GFP-positive; 75% of weakly-expressing neurons (e.g., the neuron indicated by the arrowhead) are GFP-positive (n = 213 neurons, 7 brains). Note the drivers are expressed in several additional Fmi-negative cells. Scale bar = 20 μm. (D) Immunofluorescence for Fmi and CadN on whole-mount antennal lobes of 48 h APF animals with OSN*>fmi*^*RNAi#1*^ (*peb-Gal4,UAS-Dcr-2/+;UAS-fmi*^*KK100512*^*/+*) or pan-neuronal>*fmi*^*RNAi#1*^ (*elav-Gal4,UAS-Dcr-2/+;UAS-fmi*^*KK100512*^*/+;*). In OSN*>fmi*^*RNAi#1*^ animals the remaining, relatively homogeneous glomerular signal is derived from its expression in LN (soma labeled with an arrowhead); we cannot rule out that the homogeneity is a secondary consequence of the glomerular segregation defects. Pan-neuronal>*fmi*^*RNAi#1*^ is semi-lethal, but we recovered a few adult escapers in which all Fmi immunoreactivity in the antennal lobe is lost. Scale bar = 20 μm.

**Figure S4.**
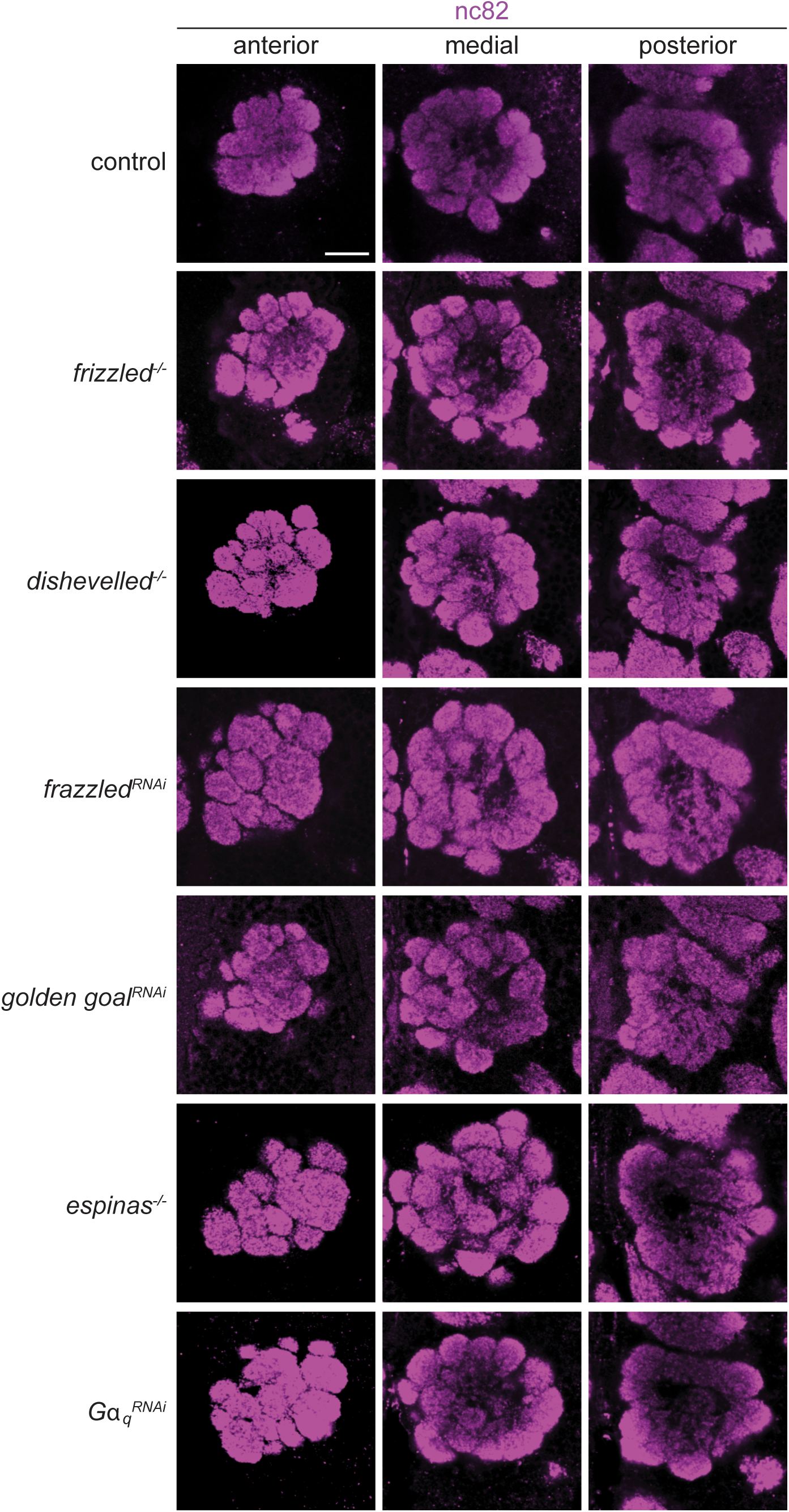
Fmi does not act with known genetic and physical interaction partners in OSNs. Representative images of nc82 immunofluorescence on whole-mount antennal lobes (three optical sections) of animals of the following genotypes (top-to-bottom; other loss-of-function alleles or RNAi lines tested, but not shown, for certain genes are also listed): control (*peb-Gal4,UAS-Dcr-2/+*), *frizzled*^*-/-*^ (*fz*^*P21*^*/fz*^*KD4A*^; also tested *fz*^*P21*^*/fz*^*H51*^), *dishevelled*^*-/-*^ (*dsh*^*1*^), *frazzled*^*RNAi*^ (*peb-Gal4,UAS-Dcr-2;Ir75b-GFP/+;fra*^*JF01457*^*/+*; also tested *fra*^*HMS01147*^ and *fra*^*JF01231*^), *golden goal*^*RNAi*^ (*peb-Gal4,UAS-Dcr-2*;*Ir75b-GFP*/*gogo*^*GD3616*^; also tested *gogo*^*HMC05937*^), *espinas*^*-/-*^ (*esn*^*KO6*^), *Gα*_*q*_^*RNAi*^ (*peb-Gal4,UAS-Dcr-2/+;Gαq*^*dsRNA*.*UAS*.*1f1*^*/+*; also tested *Gαq*^*GL01048*^ and *Gαq*^*JF01209*^). The *Ir75b*-GFP reporter, incorporated into some of these genotypes, is not shown in these images, but was indistinguishable from controls. Scale bar = 20 μm.

### Supplementary Tables

**Table S1. Chemosensory receptor genes with significant OSs in Ir neuron populations**.

Only significant OSs are listed. Values correspond to those plotted in Figure S1.

**Table S2. Set of CSSP genes**.

The gene list is from the *D. melanogaster* extracellular domain database (FlyXCDB) database^2^, with additional gene symbols appended.

**Table S3. CSSP genes with significant OSs in Ir neuron populations**.

OSs correspond to those plotted in Figure 2C.

**Table S4. CSSP genes with significant OSs unique to each population**.

**Table S5 Filtered set of transcription factors genes**.

The gene list was extracted from the FlyTF.org database^3^.

**Table S6. Transcription factor genes with significant OSs in Ir neuron populations**

OSs correspond to those plotted in Figure 2D.

**Table S7. Transcription factor genes with significant OSs unique to each population**.

**Table S8. Number of GATC motifs within candidate differentially-expressed genes contributing to significant differences**.

Values corresponds to those plotted in Figure 4D.

**Table S9.**
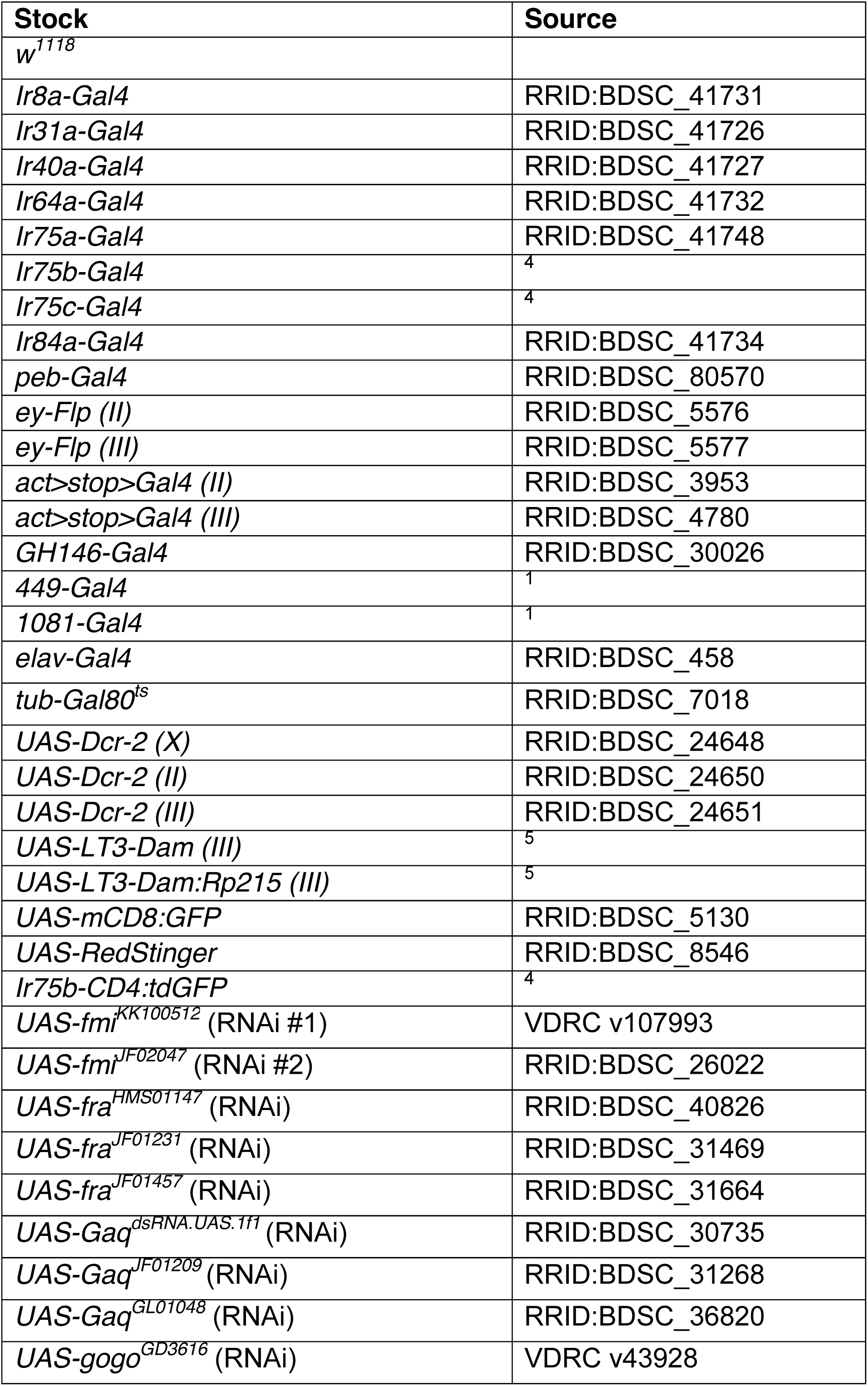
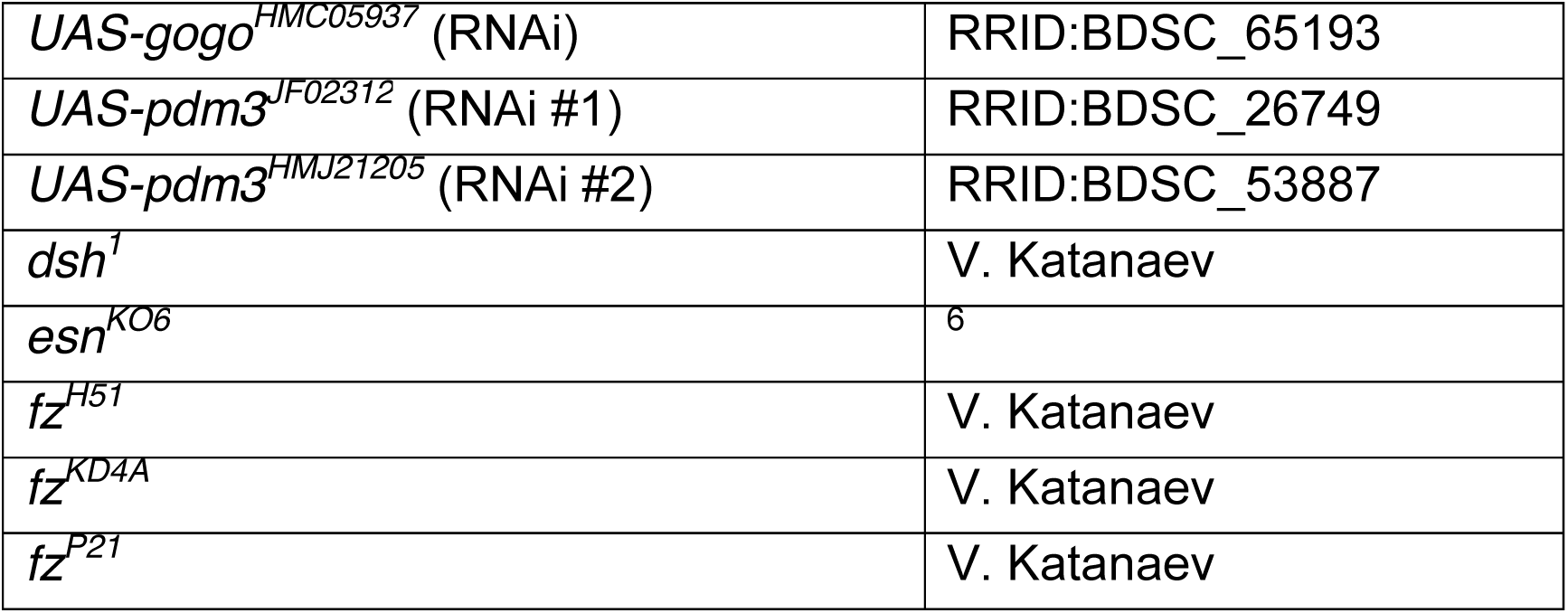
*D. melanogaster* stocks.

**Table S10.** Antibodies.

| Antibody | Dilution | Source/Reference |
| --- | --- | --- |
| Mouse mAb anti-En | 1:100 | DSHB 4D9 |
| Guinea pig anti-Pdm3 | 1:200 | <sup>7</sup> |
| Mouse mAb anti-Fmi | 1:20 | DSHB #74 |
| Guinea pig anti-IR40a | 1:200 | <sup>8</sup> |
| Rabbit anti-IR64a | 1:1000 | <sup>9</sup> |
| Rabbit anti-IR75a | 1:200 | <sup>10</sup> |
| Guinea pig anti-IR75b | 1:500 | <sup>4</sup> |
| Rabbit anti-IR75c | 1:200 | <sup>4</sup> |
| Mouse mAb anti-Bruchpilot | 1:10 | DSHB nc82 |
| Rat anti-Cadherin-N | 1:25 | DSHB Ex#8-2 |
| Rat mAb anti-Elav | 1:10 | DSHB 7E8A10 |
| Chicken anti-GFP | 1:2000 | Abcam 13970 |
| Mouse anti-GFP | 1:1000 | Invitrogen A11120 |
| Anti-RFP | 1:1000 | Abcam 62341 |
| Alexa488 anti-Guinea pig | 1:500 | Invitrogen A11073 |
| Alexa488 anti-Chicken | 1:1000 | Abcam 150169 |
| Cy3 anti-Rabbit | 1:1000 | Milan Analytica AG 111-165-1440 |
| Cy3 anti-Mouse | 1:1000 | Milan Analytica AG 115-165-166 |
| Cy5 anti-Rat | 1:250 | Jackson ImmunoResearch<br>712-175-153 |
| anti-DIG-POD | 1:300 | Roche Diagnostics 11 207 733 910 |

### Supplementary Datasets

**Supplementary Data 1. Full set of genes with significant OSs**.

**Supplementary Data 2. Candidate genes for differential occupancy based on the Likelihood Ratio Test within DESeq2**. One file contains only the list of genes, the second file contains the list of GATC fragments and the associated annotations and statistics.

**Supplementary Data 3. Candidate genes for pairwise differential occupancy based on the Wald Test with DESeq2**.

21 Bed-formatted files (representing all possible pairwise comparisons between the 7 datasets) containing statistics and annotations.

**Supplementary Data 4. Read coverage files for GATC fragments used for DESeq2 analyses**.

